# Tau-Driven Coordination of Microtubule-Actin Crosstalk in Cell-Sized Vesicles

**DOI:** 10.1101/2025.08.12.669979

**Authors:** Mousumi Akter, Jordan L. Shivers, Liya Ding, Aaron R. Dinner, Allen P. Liu

## Abstract

The coordination of microtubules (MTs) and actin filaments is essential for cytoskeletal organization, yet the factors that affect their integration remain unclear. Here, we reconstitute cytoskeletal networks in giant unilamellar vesicles to characterize MT-actin crosstalk mediated by tau, a microtubule-associated protein. We show that tau promotes the organization of MTs into diverse architectures, including bundles, clusters, and networks, depending on its concentration and vesicle size. *In vitro* assays confirm that while tau binds and bundles MTs, it does not directly bundle actin. However, tau facilitates MT-actin colocalization in the presence of actin crosslinkers with distinct properties. Fascin, which forms rigid actin bundles, significantly enhances MT-actin colocalization with tau, whereas α-actinin, which forms flexible actin bundles, induces colocalization in vesicles but not in bulk conditions. By combining cellular reconstitution and coarse-grained simulations of composite network assembly in both vesicles and bulk conditions, our findings reveal how tau-mediated cytoskeletal integration is governed by bundle mechanics and spatial confinement, providing insights into cytoskeletal organization within reconstituted synthetic cell-like systems.

## INTRODUCTION

Understanding the crosstalk between microtubules (MTs) and actin filaments is fundamental to unraveling the organizational principles of biological cells^1,2^. This crosstalk is essential for diverse cellular processes, including intracellular transport, migration, morphogenesis, and synaptic connectivity^3–6^. For instance, in neurons, the precise spatial and functional coordination between MTs and actin is critical for neurite outgrowth, synaptic plasticity, and directed cargo trafficking^7–10^. This coordination is orchestrated by a diverse array of regulatory proteins, such as microtubule-associated proteins (MAPs)^11–15^ (e.g., EB1-Shot complex, tau, kinesin-14) and actin-binding proteins (ABPs)^16–23^ (e.g., formin, profilin, cofilin), which align and stabilize cytoskeletal structures in response to both biochemical and physical cues.

Notably, certain cytoskeleton regulatory proteins possess an MT-binding domain and a proline-rich region that enables them to interact with both MTs and actin filaments. These include tau^14,15^, which is known to colocalize in axonal growth cones, especially where dynamic MTs enter actin-rich filopodia^24^. Tau misregulation disrupts cytoskeletal integration^25–27^ and is thought to play a role in various pathologies, including Alzheimer’s disease. Despite its significance, the molecular mechanisms underlying MT-actin crosstalk remain poorly understood, in part due to the difficulty in controllably probing MT-actin interactions *in vivo*.

Advances in cellular reconstitution and bottom-up synthetic biology have enabled the reconstitution of cytoskeletal systems in giant unilamellar vesicles (GUVs), providing a powerful platform to dissect the principles of cytoskeletal organization under confinement^28–32^. These systems have revealed that MTs and actin networks can each self-organize into complex structures in the presence of nucleotides, crosslinkers, and regulatory proteins^33–38^. For example, we recently showed that MT organization in GUVs is sensitive to membrane composition and nucleotide type (e.g., GTP vs. GMPCPP)^33^. Furthermore, recent work has highlighted how the identity and concentration of MAPs or ABPs, as well as the size of the encapsulating vesicle, shape emergent cytoskeletal architectures^36–39^. Encapsulation of MTs together with their associated proteins, such as MAP1 and MAP4, or motor proteins, such as kinesin, has provided valuable insights into shape changes of living cells^40,41^. Similarly, the inclusion of actin crosslinkers such as fascin and α-actinin has demonstrated that actin network architecture can be modulated so as to recapitulate structures observed *in vivo*^38,42–44^.

Despite myriad studies that reconstitute cytoskeletal features in GUVs, few have co-encapsulated MTs and actin. We propose that incorporating both MTs and actin in the same GUV compartment provides a way to probe MT-actin interactions and network formation. In particular, our study investigates MT-actin crosstalk mediated by tau, a multifunctional neuronal MT-associated protein. While tau is well known for stabilizing and bundling MTs^45–47^, its role in coordinating cytoskeletal systems under confinement has not been investigated. By combining *in vitro* cellular reconstitution with GUV encapsulation and aided by insights from coarse-grained simulations of vesicle-encapsulated and bulk composite network assembly over a broad range of conditions, we show that tau’s organizing activity is modulated by its concentration, the geometry of the vesicle, and the mechanical properties of the coexisting actin network. We demonstrate that tau concentrations and GUV diameter influence the MT network architectures. MTs colocalize with actin bundles crosslinked by fascin and not bundles crosslinked by α-actinin in bulk solution; in contrast, MTs colocalize with both types of bundles in GUVs, indicating that spatial confinement can modulate cytoskeletal integration^48,49^. We use simulations of semiflexible polymers with controlled bending rigidities to demonstrate that differences in actin bundle stiffness, together with the effects of confinement, are sufficient to account for the experimental trends^48,49^.

## RESULTS

### Tau mediates microtubule organization in confined vesicles

To examine the influence of spatial confinement on MT organization, we encapsulate MTs and tau in GUVs of varying sizes. Tau is composed of three major domains: an N-terminal projection domain, a proline-rich domain, and a C-terminal domain containing four conserved 31–32 amino acid repeat motifs^14,47^ (**Fig. 1A** top). The C-terminal domain is disordered and variably charged, which supports MT assembly and stability, while the N-terminal region of tau mediates MT interactions and is also essential for bundle organization. The repetitive units can mediate MT-binding and -stabilizing functions with tau (**Fig. 1A**, bottom).

**Figure 1.**
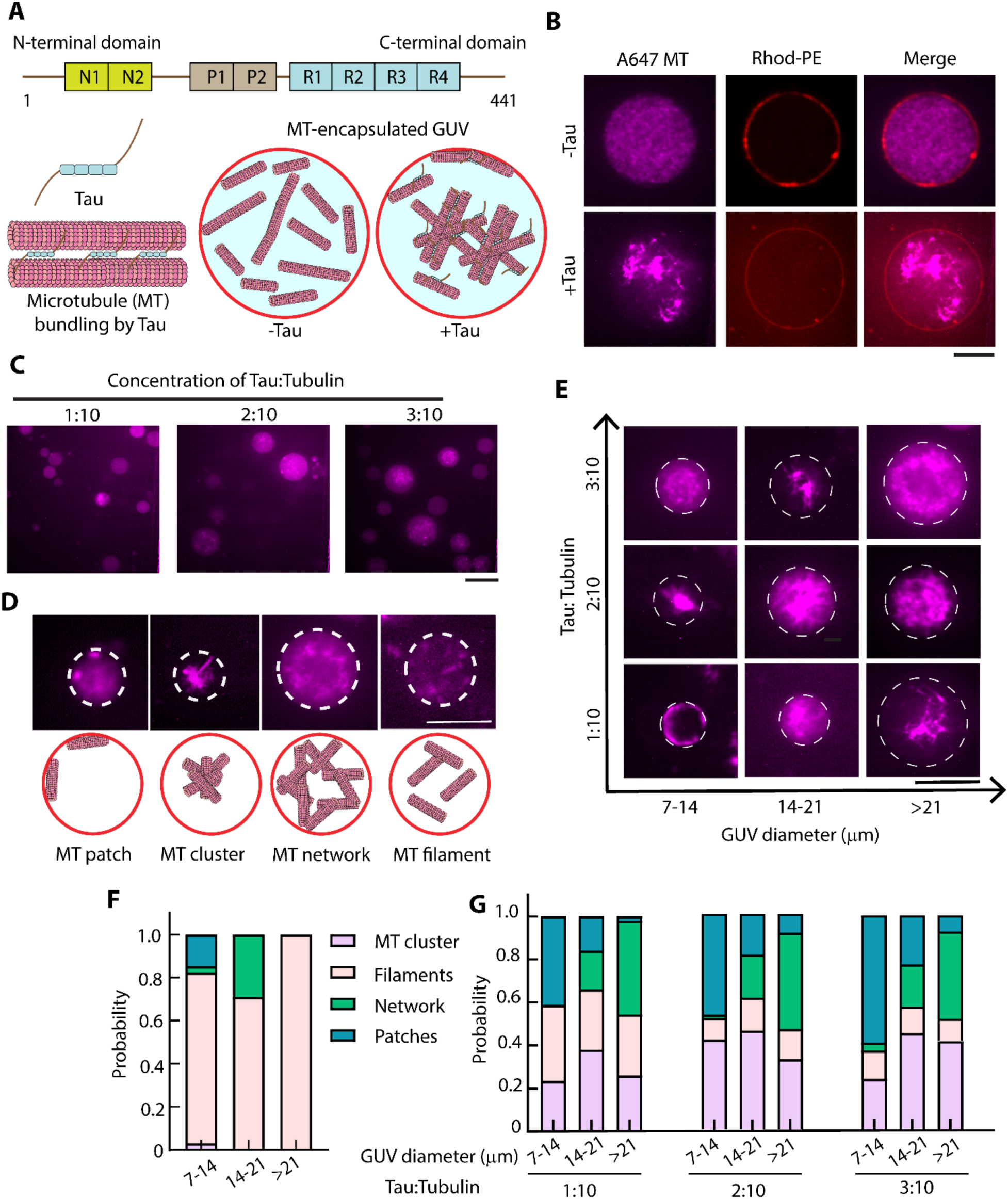
Tau-mediated organization of microtubules (MTs) in GUVs. (A) The top schematics illustrate tau protein domain structure, including the N-terminal projection domain, the proline-rich region, and the MT-binding repeat domains. The bottom schematics show a cartoon of MT bundling by tau and MT-encapsulated GUVs in the presence or absence of tau. (B) Representative confocal microscopy images of ATTO647-labeled MT-encapsulated Rhod-PE GUVs with or without tau. Scale bar: 10 μm. (C) Representative confocal microscopy images depicting various MT architectures observed at different concentration ratios of tau and tubulin. while varying tau concentration from 1 to 3 μΜ. Scale bar: 20 μm. (D) Classification of four distinct MT architectures identified in (c), along with their corresponding schematic representations. The observed structural classes are patch, filament, cluster, and network. White dashed lines outline the GUV boundaries. Scale bar: 20 μm. Tubulin concentration was kept at 10 μM in each case (total concentration in the encapsulation solution) (E) Representative confocal fluorescence images showing MT structures formed at varying tau-to-tubulin molar ratios (tubulin concentration fixed at 10 μM) across GUVs of varying diameters. Scale bar: 20 μm. (F) Cumulative probability distribution of different MT architectures in GUVs of varying sizes in absence of tau. Total 88 GUVs are considered for the analysis. (G) Cumulative probability distribution of different MT architectures in GUVs of varying sizes. The concentration ratio of tau and tubulin is indicated at the bottom of each set of bars. Number of GUVs: 208 for 1:10 (Tau:Tubulin), 480 for 2:10 (Tau:Tubulin), and 374 for 3:10 (Tau:Tubulin), totaling 1062 GUVs are considered for the analysis.

Additionally, the proline-rich region and C-terminal flanking sequences can contribute to tau’s association with both actin filaments and MTs^50,52^. To investigate tau’s role in MT-actin crosstalk and MT organization under confinement, we use full-length recombinant bovine tau containing four MT-binding repeats. We encapsulate different tau-to-tubulin concentrations in GUVs with GMPCPP, a slowly hydrolyzable analog of GTP that we previously found stabilizes reconstituted MTs^33^. The GUVs used in this study consist of 50% DOPC phospholipid and 50% PBD–PEO block copolymer, where the inclusion of PBD-PEO enhances mechanical stability and prevents vesicle rupture during MT polymerization by mitigating tubulin-induced viscosity and membrane interaction effects^51,53^.

Confocal microscopy images reveal that, in the absence of tau, tubulin encapsulated in GUVs forms a dispersed, disorganized network of MTs in GUVs (**Figure 1B**). However, the addition of tau leads to significant reorganization of MTs, which form different architectures depending on the tau-to-tubulin ratio (**Figure 1C**). Approximately 10% of GUVs contained predominantly unpolymerized tubulin, likely due to stochastic encapsulation and variations in nucleation or growth, and our analyses focused on vesicles exhibiting well-defined MT networks. At a fixed tubulin concentration (10 μM), increasing tau concentration from 1 to 3 μM leads to the emergence of four major classes of MT structures: patch, filament, cluster, and network. These structures are schematically represented in **Figure 1D**. In the patch class, GUVs contain one or more patches of short, bundled MTs, on the membrane. As MTs polymerize and form bundles in the presence of tau, the longer MT bundles can undergo buckling near the vesicle membrane due to steric confinement and non-specific interactions between MTs and the membrane, which may promote the formation of MT patches within GUVs. For the remainder of the classes, the MTs are primarily in the lumen. The filament class features MT filaments dispersed throughout the GUV lumen, exhibiting lower fluorescence intensity compared to the patch structures. In the cluster class, MTs are extensively crosslinked, forming dense, compact architectures in the lumen, while in the network class, the MTs are interconnected but dispersed compared with the cluster structures. We also note that the observed MT patch formation is consistent with results obtained using DOPC-only GUVs. Experiments performed in the absence of PBD–PEO incorporation exhibited similar MT patch structures, indicating that the inclusion of PBD–PEO does not qualitatively influence the underlying cytoskeletal organization (**Figure S1**).

To classify MT-encapsulated GUVs based on the above phenotypes, we adopt a supervised machine learning approach (*See Supporting Information*). MT structures encapsulated in GUVs are segmented from microscopy images using preprocessing, circular detection, and intensity-based filtering to isolate relevant regions (**Figure S2**). To ensure consistency across a large dataset and enable quantitative comparison, MT organization was analyzed at the equatorial plane, although this approach may omit structural information from other focal planes. A subset of image patches is manually annotated into four structural categories and used to fine-tune an EfficientNet-B0 convolutional neural network for automated classification^54^. The trained model is used to classify the full dataset, and structural distributions are analyzed across varying tau concentrations and GUV size ranges to study MT organization dynamics (**Figure S3**). **Figures 1E-1G** show that the architecture of MT assemblies is influenced by both the tau concentration and GUV size. In the absence of tau, MTs predominantly form dispersed filamentous structures inside GUVs of different sizes (**Figure 1F**). At the lowest tau concentration (1:10), the network class dominates in larger GUVs, while smaller GUVs show more of the patch class. As the concentration increases, the cluster class becomes more prominent, particularly in smaller and intermediate-sized GUVs.

### Tau facilitates MT-actin crosstalk in solution

Having shown that tau can organize MTs in GUVs, we next investigate the interactions between tau, MTs, and actin filaments. We start by characterizing MT-tau interactions *in vitro*, prior to exploring tau-mediated MT–actin crosstalk. TIRF microscopy images show that, in the presence of 2 μM tau, MTs form organized, bundle-like structures, which are not observed in the absence of tau (**Figure 2A**). However, no actin bundling is observed under the same conditions (**Figure S4**). These observations are consistent with previous reports^46,55,56^ and further supported by results from an MT-actin-tau co-pelleting assay (**Figure S5**). The assay is performed with a total volume of 40 µL, followed by ultracentrifugation and separation into supernatant (20 µL) and pellet (20 µL) fractions (see Methods). As expected, preformed MTs incubated with tau are enriched in the pellet, consistent with tau’s known binding to MTs. In contrast, tau incubated with actin alone does not result in detectable actin in the pellet, confirming that tau does not bind actin directly. Unexpectedly, a fraction of F-actin is detected in the pellet when co-incubated with both MTs and tau, which may result from weak associations among the proteins. We image co-polymerization of MTs (10 μM) and actin (5.5 μM) with tau by TIRF microscopy. We observe MT bundles along with actin filaments, suggesting that tau mediates MT but not actin bundling (**Figure 2B**). This is also consistent with previously reported observations of tau-mediated MT-actin crosstalk in bulk^14,15,57^.

**Figure 2.**
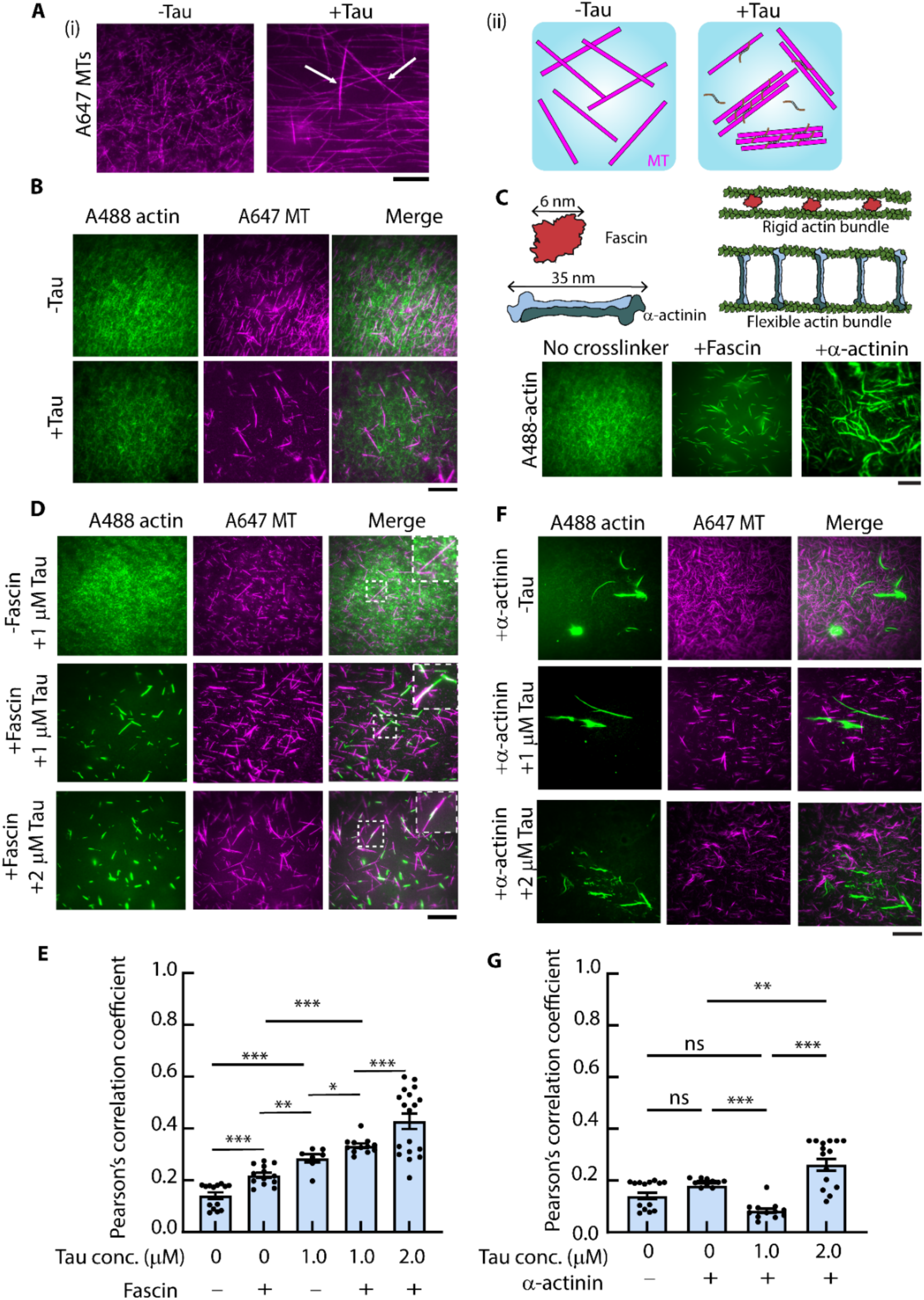
Tau-mediated MT-actin crosstalk in vitro. (A) (i) Representative TIRF microscopy images depicting MT organization in the presence or absence of 2 µM tau (ii)The schematic illustrates MT organization in the presence or absence of tau in bulk solution. (B) Representative TIRF microscopy images comparing the organization of 10 µM MTs and 5.5 µM actin filaments in the presence and absence of 1 µM tau after 1 h of polymerization at 37 °C. Tau to tubulin ratio is 1:10. (C) The upper panel illustrates the mechanisms of actin crosslinking by actin-bundling proteins fascin and α-actinin. The lower panel shows representative TIRF microscopy images demonstrating rigid actin bundles induced by fascin and flexible actin bundles formed by 1.2 µM α-actinin. (D) Representative TIRF microscopy images showing 10 µM MTs and 5.5 µM actin filaments in the presence or absence of tau and the actin crosslinker fascin (1.2 µM). The white dashed box highlights a magnified region showing colocalized MT-actin structures. The tau-to-tubulin ratio in each condition is 1:10, 1:10, and 2:10, respectively (top to bottom). (E) Comparison of the effects of varying tau concentrations on MT-actin colocalization in solution in the presence or absence of 1.2 µM of fascin, as measured by Pearson’s correlation coefficient. (F) Representative TIRF microscopy images showing MTs and actin filaments in the presence or absence of tau and α-actinin (1.2 µM). The tau-to-tubulin ratio in each condition is 1:10, 1:10, and 2:10, respectively (top to bottom). (G) Comparison of the effects of tau on MT-actin colocalization in solution in the presence or absence of α-actinin (1.2 µM), as measured by Pearson’s correlation coefficient. The tau concentration was 1.0 µM. * represents p < 0.05, *** represents p < 0.001, and ns represents non-significant. Error bars represent the SEM. A minimum of 10 ROIs was considered for the analysis from three independent experiments. Scale bar: 10 μm.

Motivated by the observation of MTs in actin-rich neuronal projections in the presence of tau^24^, we introduce two actin bundling proteins, fascin and α-actinin, with distinct biophysical properties. Fascin, a short (6 nm) crosslinker, induces the formation of rigid, parallel actin bundles, whereas α-actinin, a long (35 nm) crosslinker, promotes the formation of loosely organized, flexible bundles (**Figure 2C**). We estimate the persistence length, a measure of filament rigidity, of tau-mediated MT bundles as well as fascin- and α-actinin-mediated actin bundles by measuring the end-to-end distance and contour length of individual bundles and fitting the data to Equation (1) (see Methods). For persistence length measurements, only isolated segments of actin or MT bundles are analyzed. As shown in **Figure S6**, the average persistence length of tau-induced MT bundles is 475.7±8.9 µm, consistent with previous reports^58^. Fascin-induced actin bundles exhibit an average persistence length of 902.8±27.0 µm, while α-actinin-induced bundles are more flexible, with an average persistence length of 285.3±6.3 µm. For comparison, actin filaments have a persistence length of ∼15 µm^59^. These results confirm that fascin-crosslinked actin bundles are more rigid than those formed by α-actinin.

Interestingly, when tau is co-incubated with the short crosslinker fascin, a significant increase in MT-actin colocalization is observed, as evident from the colocalized filamentous structures shown in **Figure 2D** and the comparison of Pearson’s correlation coefficients between the tubulin and actin intensities in **Figure 2E**. In contrast, with α-actinin, tau does not promote significant MT-actin colocalization (**Figure 2F-G**), indicating that the type of actin crosslinker can affect the extent of cytoskeletal crosstalk in solution. To evaluate the effect of crowding factor on MT–actin association, we performed additional experiments in the presence of 0.2% methylcellulose and observed no significant change in the colocalization ratio compared to the non-crowded condition (**Figure S7**).

### Actin crosslinkers mediate cytoskeletal organization in confined GUVs

We next examine whether tau facilitates MT-actin crosstalk in confined GUV environments in the presence of fascin. We polymerize MTs and actin filaments with 0 to 3 μM tau in the presence of 1.2 μM of fascin in GUVs. Confocal microscopy images show colocalization of MTs and fascin-bundled actin filaments in GUVs in the presence of tau (**Figure 3A**). Corresponding fluorescence intensity profiles in **Figure 3B** illustrate the spatial distribution of MTs and actin filaments in the GUV interior, showing overlapping MT and actin intensities. We quantify colocalization using Pearson’s correlation coefficient for the tubulin and actin intensities (**Figure 3C**), and this analysis reveals significantly more colocalization in the presence of tau (**Figure 3D**). In the absence of tau, the Pearson’s coefficient is largely independent of GUV size, whereas increasing tau concentration enhances overall colocalization without a clear size dependence, indicating that tau strengthens MT–actin interactions without altering their size-dependent organization (**Figure 3E**). These results suggest that tau, together with fascin-induced actin bundles, promotes cytoskeletal integration of MTs and actin under confined conditions. We also confirmed that differences in MT-actin colocalization between in solution and GUV were not due to imaging modality or resolution, as Pearson’s correlation coefficients were consistent when TIRF images were downsampled to match confocal resolution under identical experimental conditions (**Figure S8**).

**Figure 3.**
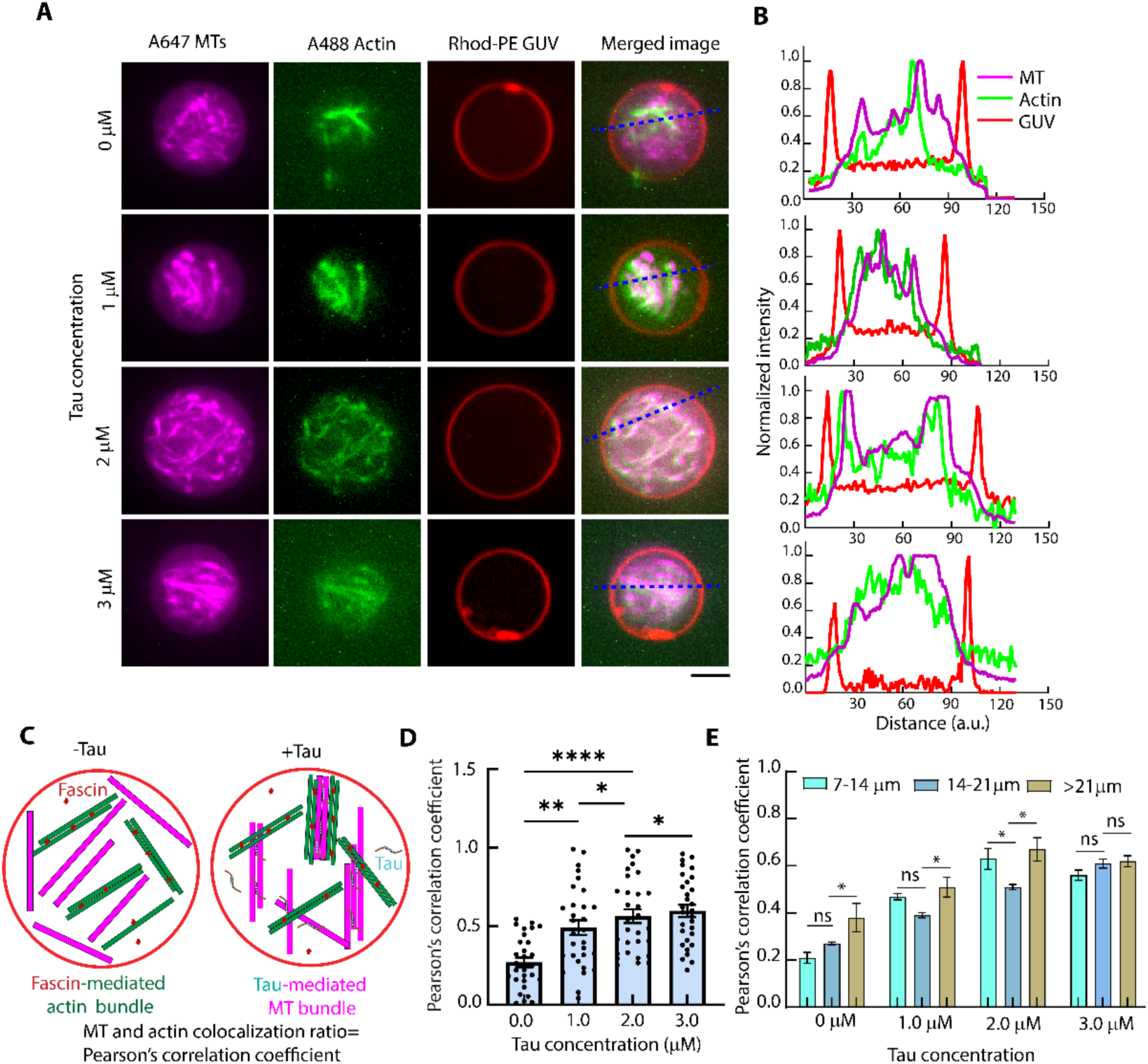
Tau-mediated MT-actin crosstalk in GUV in the presence of fascin. (A) Representative confocal microscopy images showing tau-mediated MT-actin colocalization in Rhod-PE GUV in the presence of 1.2 µM fascin. These images represent single z-slices acquired at the equatorial plane of the GUVs. Scale bar: 10 μm. (B) Normalized fluorescence intensity profile along the blue line in the merged channel of panel (A), illustrating the spatial distribution of MTs and actin filaments in the GUV. The magenta, green, and red lines represent fluorescence intensities of MT, actin, and GUV membrane, respectively. (C) Schematic illustrations depicting MT and fascin-bundled actin organization in GUVs, both in the presence and absence of tau. (D) Comparison of the effects of varying tau concentrations on MT-actin colocalization in GUVs, as measured by Pearson’s correlation coefficient for the tubulin and actin intensities. **** represents *p* < 0.0001, *** represents *p* < 0.001 * *p* < 0.05 and ns = non-significant. Error bars represent SEM. The scatter plots display individual data points from ∼30 GUVs, and SEM is calculated across these 30 GUVs from three independent experiments. (E) Dependence of MT–actin colocalization on GUV size at different concentrations of tau, quantified using Pearson’s correlation coefficient between MT and actin fluorescence intensities at a fixed fascin concentration of 1.2 µM. Error bars represent the SEM. Here, ns = non-significant and * *p* < 0.05. For each condition, more than 30 GUVs from three independent experiments were analyzed.

To determine the effects of the actin crosslinker on MT-actin organization in GUVs, we vary the concentrations of fascin or α-actinin at 2 μM of tau. Confocal microscopy images and colocalization analysis show that MT-actin networks colocalize in GUVs with increasing fascin concentrations (**Figure 4A, B**). Strikingly, MT-actin networks also colocalize in GUVs with increasing α-actinin concentrations (**Figure 4D, E**), in contrast to MT-actin networks in bulk solution (**Figure 2E**). These results suggest that both the type of actin crosslinker and spatial confinement critically influence the extent of MT-actin colocalization. Figures 3D and 4B correspond to independent experimental series conducted on different days, one performed with 1.2 μM fascin and varying tau concentrations (yielding ∼50% colocalization) and the other with 2.0 μM tau and varying fascin concentrations (yielding ∼35% colocalization). Such variations in absolute colocalization values likely arise from day-to-day experimental differences; however, both datasets consistently demonstrate enhanced MT-actin colocalization in the presence of fascin and tau.

**Figure 4.**
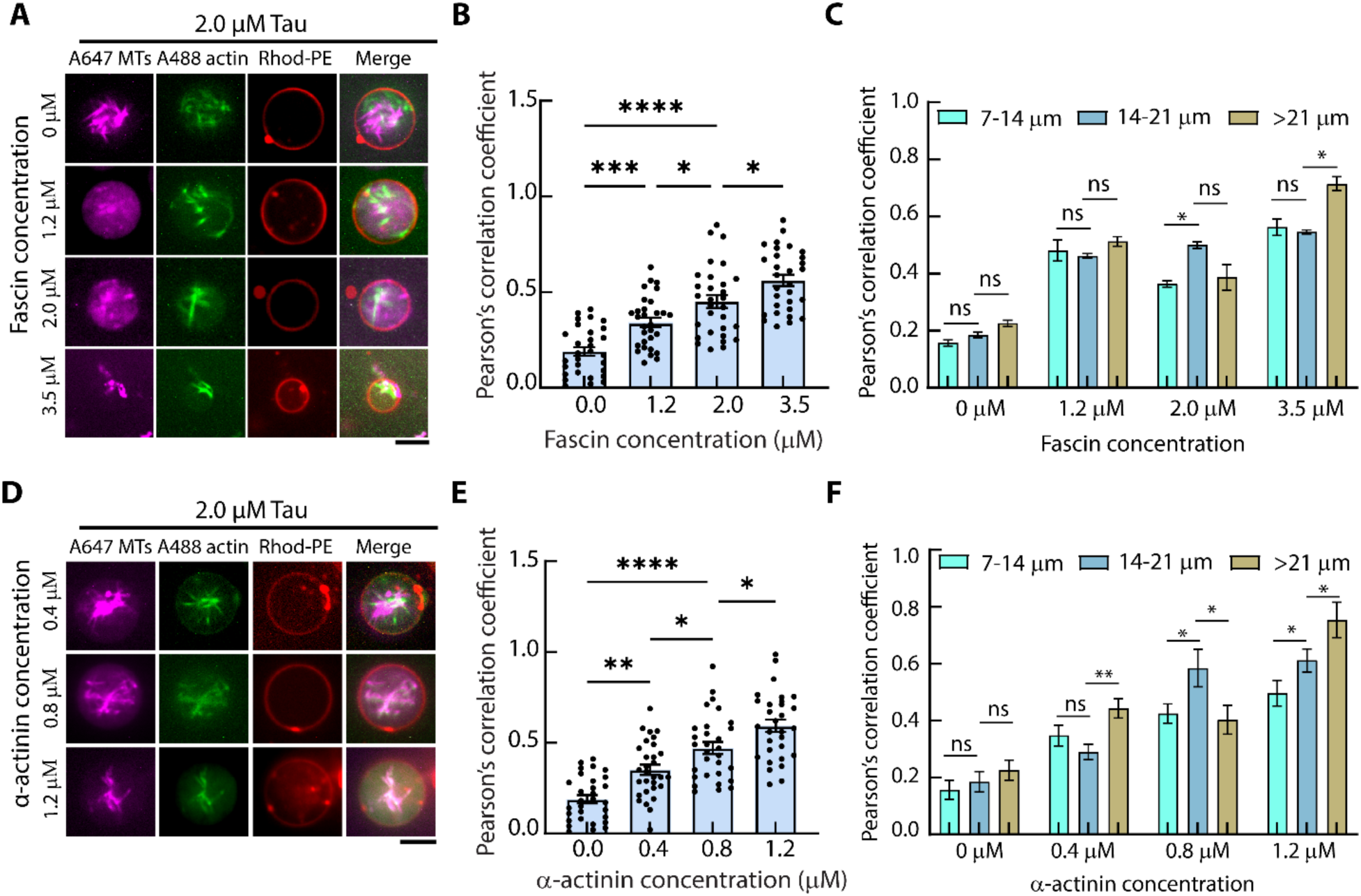
Tau-mediated MT-actin crosstalk in GUVs in the presence of fascin or α-actinin. (A) Representative confocal fluorescence images show MT and actin organization in Rhod-PE- GUVs in the presence of 2 µM tau and different concentrations of fascin ranging from 1.2 μM to 3.5 μM. Scale bar: 20 μm. These images represent single z-slices acquired at the equatorial plane of the GUVs. (B) Quantitative analysis of MT-actin colocalization using Pearson’s correlation coefficient at varying fascin concentrations. Increased fascin concentrations correlate with a statistically significant increase in MT-actin colocalization. (C) Dependence of MT–actin colocalization on GUV size as a function of fascin concentration, quantified by the Pearson’s correlation coefficient between MT and actin fluorescence intensities at a fixed Tau concentration of 2.0 µM. Error bars represent the SEM. For each condition, more than 30 GUVs from three independent experiments were analyzed. (D) Representative confocal fluorescence images show MT and actin organization in Rhod-PE GUVs in the presence of 2 µM tau and different concentrations of α-actinin ranging from 0.4 to 1.2 μM (single z-slice). Scale bar: 20 μm. (E) Quantitative analysis of MT-actin colocalization using Pearson’s correlation coefficient at varying α-actinin concentrations. (F) Variation of Pearson’s correlation coefficient with α-actinin concentration at a fixed Tau concentration of 2.0 µM. Error bars represent the SEM. For each condition, more than 30 GUVs from three independent experiments were analyzed. ns = non-significant, * *p* < 0.05, ** *p* < 0.01, *** *p* < 0.001, and **** *p* < 0.0001. Error bars represent the SEM (n > 50 GUVs, five experiments for B and E).

Figures 4C and 4F illustrate the dependence of MT–actin colocalization on vesicle size at varying concentrations of fascin or α-actinin. In the absence of actin crosslinkers, MT–actin colocalization remains largely independent of vesicle size. At 1.2 µM fascin, the Pearson’s coefficient is approximately 0.5 across all GUV sizes. Increasing the fascin concentration to 2.0 µM results in a peak colocalization (∼0.5) in moderately sized vesicles, whereas at 3.5 µM fascin, the coefficient rises to 0.6–0.7 and reaches its maximum in larger vesicles. Similarly, at 1.2 µM α-actinin, the Pearson’s coefficient remains moderate in smaller GUVs (7–14 µm) but increases markedly to ∼0.8 in larger vesicles (>21 µm), indicating that both α-actinin concentration and compartment size synergistically enhance cytoskeletal organization and MT–actin colocalization within GUVs.

To directly assess the relationship between MT architecture and actin–MT organization, we analyze MT structural patterns and their degree of colocalization with actin across varying tau:tubulin and actin crosslinker ratios. As shown in **Figure S9**, in the absence of actin crosslinkers, filamentous MTs predominate, and actin–MT colocalization remains low. The addition of actin and fascin markedly increases the occurrence of network-like MT architectures, particularly in larger GUVs (>21 µm), which coincides with a pronounced enhancement in actin–MT colocalization. At higher tau:tubulin ratios (1:10–3:10), MT networks become progressively dominant while patches decrease, and clusters persist across all GUV sizes. The presence of fascin or α-actinin further promotes MT network formation and enhances actin–MT colocalization, suggesting that increased actin bundle stiffness facilitates the spatial organization of MTs into interconnected networks. Our results reveal a positive correlation between the probability of network formation and the Pearson’s correlation coefficient of actin–MT fluorescence intensities, indicating that higher degrees of MT networking are associated with stronger actin–MT colocalization. Collectively, these findings demonstrate that GUV size, tau concentration, and actin crosslinker composition synergistically govern MT architectural transitions and the extent of actin–MT colocalization, with stronger actin crosslinking promoting the emergence of robust, highly interconnected MT–actin networks.

### Model for tau-mediated MT-actin crosstalk

Experimentally, we observe colocalization of MTs with fascin-bundled actin filaments but not α-actinin-bundled filaments in bulk solution, as well as colocalization in GUVs with α-actinin-bundled filaments. We propose that the differences in colocalization patterns derive from the marked differences in bending rigidities of the actin bundles formed by these crosslinkers: fascin produces tightly packed, rigid bundles, whereas α-actinin produces more flexible, loose actin bundles (Figure 5A).

**Figure 5.**
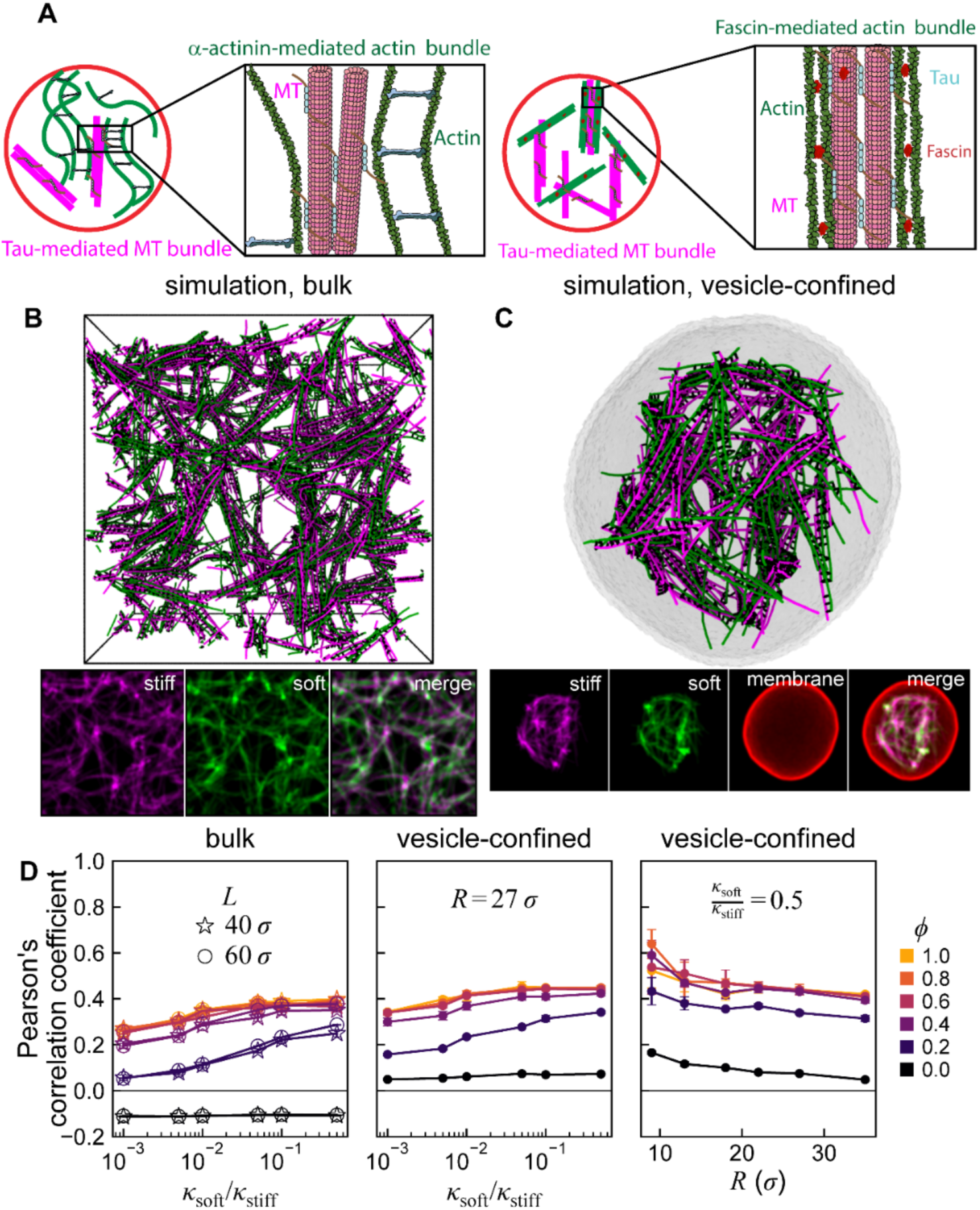
Stiffness compatibility and confinement enhance MT-actin colocalization. (A) Schematic overview of tau-mediated interactions between MTs and actin filaments in the presence of α-actinin and fascin in a GUV. The magnified views depict how tau both bundles MTs and bridges MTs and actin filaments, while α-actinin and fascin produce actin bundles with distinct properties. Bundles crosslinked by α-actinin, a long and flexible actin-crosslinking protein, are looser and more flexible than the tight, rigid bundles produced by the shorter fascin molecules. In the presence of tau, stiffer fascin-linked actin bundles readily colocalize with MTs both in GUVs and in bulk solution, whereas more flexible α-actinin-linked actin bundles only colocalize with MTs in GUVs but not in bulk solution. To understand these observations, we perform minimal coarse-grained simulations of the assembly of crosslinked networks of two semiflexible filament species, representing MTs with a “stiff” species of fixed bending rigidity κ_stiff_ and representing actin bundles with a more flexible “soft” species with a variable bending rigidity κ_soft_, in either (B) bulk conditions, with periodic boundaries, or (C) confined in flexible fluid vesicles. For a given bending rigidity ratio κ_soft_/κ_stiff_ and crosslinker coverage fraction ϕ, we allow the networks to assemble for a fixed duration and generate artificial fluorescence images of the final assembled simulation configurations for each particle type, shown beneath the simulation configurations in (B) and (C). Using these images, we quantify the colocalization between the stiff and soft polymer types by the Pearson’s correlation coefficient *r* between their respective fluorescence intensities. (D) Increasing either the bending rigidity ratio κ_soft_/κ_stiff_ or the crosslinker coverage fraction ϕ results in an increase in colocalization both (left) in bulk networks and (center) in networks confined in vesicles of radius *R*; (right) reducing the vesicle radius *R* leads to increased colocalization between the two species.

To explore this hypothesis, we perform coarse-grained simulations of the assembly of crosslinked semiflexible polymer networks composed of two filament types: a more flexible species representing actin bundles and a stiffer species representing MTs (**Figure S10, S11**). By varying the bending rigidity *κ*_soft_ of the more flexible species, we shift between a regime representative of flexible, α-actinin-crosslinked actin bundles and a regime representative of rigid, fascin-crosslinked actin bundles. We note that the simulations explore a broader range of bending rigidities than observed experimentally, in order to systematically map colocalization trends and confirm their robustness. To model the influence of tau concentration, we decorate filaments with a specified coverage fraction *ϕ* of uniformly distributed “sticky” sites capable of forming reversible crosslinks with other nearby filaments. To capture GUV confinement, vesicles are modeled using a coarse-grained particle-based membrane model^60^ (**Figure S12**). After an initial equilibration stage, we activate crosslinking and allow simulations to proceed for a fixed duration, after which the final network structure is analyzed. To facilitate direct comparison with experimental image analysis, we post-process the final simulation configurations to generate artificial fluorescence intensity maps (**Figure S13, S14**). Colocalization between stiff and soft filaments is then quantified using Pearson’s correlation coefficient, mirroring the experimental image analysis protocol. Further details about the simulation methods and associated colocalization analysis are provided in *Methods* and in *Supplementary Information*.

In Figure 5B**, C,** and **Figure S15-S24**, we show representative configurations for bulk and vesicle-confined networks. In the left panel of Figure 5D and **Figure S25**, we plot the computed Pearson’s correlation coefficient for bulk systems as a function of the bending stiffness ratio *κ*_soft_/*κ*_stiff_, for varied crosslinker (sticky site) coverage fraction *ϕ*. For all nonzero *ϕ*, increasing the bending stiffness ratio *κ*_soft_/*κ*_stiff_ leads to an increase in Pearson’s correlation coefficient, indicating increased colocalization. These results indicate that changes in actin bundle stiffness are sufficient to account for the observed increases in colocalization at higher concentrations of fascin and α-actinin; that is, the mechanism is generic and does not rely on specific molecular details. Moreover, our simulations show that increasing *ϕ*, an analog of the tau concentration, generically leads to increased colocalization between the two species. In **Figure S26**, we show that this behavior persists even as the filament length *L*_*α*_is varied. We note that the two bulk system sizes tested (periodic box side lengths *L*_box_ = 40*σ* and 60*σ*, where *σ* is a unit length) yield consistent values of Pearson’s correlation coefficient across *ϕ* and *κ*_soft_/*κ*_stiff_, suggesting that our bulk results are insensitive to system size. To confirm that the simulations reached steady state, we quantified the time evolution of the average number of crosslinks per filament for different coverage fractions (*ɸ*), which plateaued after approximately 1 × 10⁶ time steps, confirming that the chosen simulation duration (1.5 × 10⁶ time steps) is sufficient for network stabilization (**Fig. S27**).

Our simulations furthermore recapitulate the experimental observation that confinement leads to increased MT-actin colocalization in the presence of tau. In the center panel of Figure 5D, we show that, in vesicles of fixed size, the Pearson’s correlation coefficient increases relative to the bulk case under all conditions but exhibits the same tendency to increase with both the crosslinker coverage fraction *ϕ* and polymer bending stiffness ratio *κ*_soft_/*κ*_stiff_. We likewise consider the influence of varying the vesicle radius *R* for networks with fixed bending stiffness ratio *κ*_soft_/*κ*_stiff_ = 0.5 (Figure 5D, right panel). We find that, for all values of the crosslinker coverage fraction *ϕ*, increasing confinement (decreasing vesicle radius *R*) increases the colocalization between the soft and stiff filaments. This may explain our experimental observation that more flexible α-actinin-bundled actin colocalizes with MTs in GUVs but not in bulk. Put together, our experiments and simulations demonstrate the importance of confinement, crosslinker properties, and tau concentration in modulating the emergent cytoskeletal organization in synthetic cellular environments.

## DISCUSSION

Co-polymerization of MTs and actin filaments in a single system poses inherent challenges due to their distinct assembly dynamics and buffer requirements. Previous studies have explored MT-actin composites in bulk solution, particularly in relation to contractility under different crosslinker or motor protein conditions^61–64^. A recent protocol by Vendel *et al.* reconstituted MT-actin networks within water-in-oil emulsion droplets using preassembled actin filaments and exogenous linkers^65^. In contrast, our GUV system enables simultaneous co-polymerization of actin and MTs within a membrane-bound environment, providing a more physiologically relevant platform to study cytoskeletal organization under confinement. To our knowledge, we present the first cellular reconstitution of MTs and actin filaments polymerized together in GUVs, allowing us to investigate how spatial confinement and tau-mediated MT bundling influence MT-actin network architecture. This confined system reveals unique features of cytoskeletal organization not accessible in unconfined environments and provides a new platform for dissecting MAP-dependent cytoskeletal interactions.

Our results reveal that cytoskeletal filaments’ mechanical properties, crosslinker mechanics, and spatial confinement together govern the patterning of the cytoskeleton. Fascin and α-actinin, two well-characterized actin-bundling proteins, exhibit distinct bundling behaviors. As previously shown by Kovar and coworkers, these proteins self-sort in mixed systems: each promotes the formation of its preferred bundle type while excluding the other^66^. Fascin forms tightly packed, rigid bundles with narrow filament spacing, whereas α-actinin produces looser, more compliant bundles with wider spacing between filaments. We observed a comparable phenomenon in our mixed MT-actin system. When actin was bundled by fascin, it formed dense cables that readily aligned and colocalized with tau-bundled MTs in bulk solution, suggesting structural compatibility between the rigid actin bundles and the thick MT arrays. In contrast, α-actinin produced flexible, long actin bundles that failed to co-localize efficiently with the tau-MT bundles in bulk solution. These findings establish a mechanistic link between MT architecture and actin–MT colocalization. Fascin- or α-actinin-mediated actin bundling not only modulates actin stiffness but also provides a physical scaffold that promotes lateral MT alignment and crosslinking, thereby enhancing MT network formation and reinforcing cytoskeletal integration within GUVs. These observations reinforce the notion that the structural and mechanical characteristics of the actin network, imparted by its crosslinkers, play a central role in determining how well it can integrate with tau-organized MT networks. While previous studies showed that actin crosslinkers such as fascin and α-actinin sort into distinct domains in actin networks^38,66^, our work demonstrates that the stiffness of actin bundles can influence crosstalk in a mixed cytoskeletal system. Although we did not directly observe differential association with MTs, our findings suggest that crosslinker mechanics and filament context together modulate network architecture, consistent with other reports^67,68^. In our experiments, bundle persistence lengths (300-900 µm) greatly exceed vesicle radii (5-25 µm), such that boundary exclusion effects likely concentrate filaments away from the membrane while restricting their orientational degrees of freedom, enhancing the probability of crosslink formation compared to bulk conditions.

An essential feature of our experimental system is the spatial confinement imposed by the GUV membranes. Kandiyoth and Michelot have argued that encapsulating actin networks in cell-sized volumes, compared to bulk reconstitution, fundamentally changes the principles of self-organization by introducing limited monomer pools, restricted diffusion, and geometric constraints^69^. Competition for actin monomers among different actin structures is amplified in the closed environment of the vesicle. Encapsulation leads to resource-sharing effects and concentration gradients that do not arise in open systems^38^. In this context, the observed spatial organization and differences in filament network compactness may be influenced by the available cytoskeletal components in the limited volume of the vesicle. While GUV size couples geometric confinement with the total encapsulated protein amount, our analysis using normalized parameters indicates that the tau-to-tubulin ratio, rather than vesicle diameter alone, also governs MT architecture.

More broadly, vesicle geometry likely influences cytoskeletal patterning by modulating filament orientation and interaction. In reconstituted systems, curved membranes have been shown to affect cytoskeletal organization by constraining filament alignment and promoting spatial partitioning^34,70,71^. Our findings demonstrate that even minimal reconstituted systems can recapitulate key aspects of cytoskeletal patterning observed in cells. The emergent colocalization and segregation of MTs and actin in both bulk and encapsulated environments reflect how cells employ MAPs and ABPs to define functionally distinct cytoskeletal domains. For example, in neurons, tau organizes MTs into parallel bundles in axons, while actin, influenced by bundlers like fascin or α-actinin, contributes to the formation of dynamic protrusions such as growth cones and dendritic spines. Our findings substantiate this functional segregation: tau mediates MT bundling, and the identity of the actin crosslinker dictates how actin filaments associate with the MT scaffold.

Our simulations suggest that these behaviors arise from general physical principles rather than relying on molecular details. Specifically, we find that, for a system of two semiflexible filament species—a stiff species with large bending rigidity *κ*_stiff_ and a softer species with *κ*_soft_ < *κ*_stiff_—in the presence of a crosslinker (representing tau), colocalization between the two species increases with the stiffness of the softer species, the crosslinker concentration, and the degree of confinement. This may offer insight into our experimental observation of stronger colocalization between α-actinin-bundled actin and MTs in GUVs than in bulk. Visual inspection confirms that this increased colocalization coincides with the formation of aligned, crosslinked filament bundles containing both species. The effects of stiffness, crosslinker concentration, and confinement likely reflect both thermodynamic and kinetic contributions. Bundling suppresses bending fluctuations, incurring an entropic cost that scales with filament bending rigidity *κ* as *κ*^-1/4/ 72,73^; increasing the stiffness of the soft species thus reduces this cost, facilitating bundle formation. However, since this entropic cost scales weakly with bending stiffness, we cannot rule out a possible influence of kinetic effects on colocalization. Kinetically, more flexible filaments are more likely to produce entanglements and sterically hinder rearrangement, slowing the formation of ordered bundles. Reversible crosslinking introduces constraints that both disproportionally reduce the entropy of more flexible filaments and increase the timescales of reorganization of disordered configurations. Confinement reduces the accessible configurational space, lowering the entropic penalty of bundling^74,75^, and increases the frequency of filament-filament collisions, effectively raising local concentrations of crosslinkers and filaments and thus increasing the on-rate for crosslink formation^76,77^. While our simulation results do suggest that differences in bundle stiffness could explain the observed difference in colocalization between fascin- and α-actinin-bundled actin, they do not rule out other possible mechanisms, such as a difference in tau-actin interaction strengths in the case of fascin- vs α-actinin-bound actin.

Finally, this work highlights the utility of cellular reconstitution approaches for dissecting the fundamental principles of cytoskeletal organization. By precisely controlling molecular composition, geometry, and confinement, we gain insight into how cells might regulate their internal architecture through physical and biochemical cues. Future studies could explore how varying vesicle size, membrane curvature, or incorporating additional MAPs and motor proteins influence spatial patterning. Altogether, our findings reveal that tau-driven MT-actin crosstalk in cytoskeletal organization in confined environments is dependent on actin bundling and suggest that filament mechanics and spatial constraints are equally critical for understanding cytoskeletal behavior in synthetic cell-like systems.

## METHODS

### Proteins and reagents

Unlabeled cycled tubulin was obtained from PurSolutions LLC or generously provided by Puck Ohi (University of Michigan). HiLyte 647 porcine tubulin, unlabeled actin, ATTO 488-labeled actin, and porcine brain tau were purchased from Cytoskeleton Inc. (USA). α-Actinin used in the experiments was kindly provided by Dr. David Kovar (University of Chicago).

Fascin was expressed in *Escherichia coli* as a glutathione-S-transferase (GST) fusion protein and purified following established protocols^78^. Specifically, *E. coli* BL21(DE3) cells were transformed with a pGEX-6P-1 plasmid encoding fascin fused to GST via a PreScission protease cleavage site. Transformed cells were cultured at 37 °C with shaking at 220 rpm until the optical density at 600 nm reached 0.5–0.6. Protein expression was induced by adding 0.1 mM isopropyl β-D-1-thiogalactopyranoside (IPTG), and cells were incubated at 24 °C for 8 hours. Following induction, cells were harvested by centrifugation at 4,000 × *g* for 30 minutes, washed once with phosphate-buffered saline (PBS), and resuspended in lysis buffer (20 mM K-HEPES, pH 7.5, 100 mM NaCl, 1 mM EDTA, and 1 mM phenylmethylsulfonyl fluoride (PMSF)). Cell lysis was performed by sonication, and the lysate was subsequently centrifuged at 45,000 × *g* for 25 minutes. The resulting supernatant was loaded onto a 1 mL GSTrap FF affinity column (Cytiva) using an ÄKTA Start fast protein liquid chromatography system. The column was washed with 15 mL of washing buffer (20 mM K-HEPES, pH 7.5, 500 mM NaCl), followed by incubation with 2 mL of cleavage buffer (50 mM Tris-HCl, pH 7.5, 150 mM NaCl, 1 mM EDTA, and 1 mM dithiothreitol) containing 50 units of PreScission protease (GenScript) at 4 °C overnight to remove the GST tag. GST-free fascin was eluted using 5 mL of PBS, followed by dialysis against 1 L of PBS twice for 3 hours and once overnight at 4°C. Protein concentration was determined using a NanoDrop spectrophotometer, and samples were concentrated using a Centricon filter (Millipore). The purified fascin was aliquoted and stored at −80 °C.

### *In vitro* MT*- and* actin-binding assay

Taxol-stabilized GMPCPP MTs were prepared by polymerizing 10 µM tubulin at 37 °C in BRB80 buffer (80 mM PIPES, 1 mM EGTA, 1 mM MgCl₂; pH ∼6.8) in the presence of 1 mM guanylyl-(α,β)-methylenediphosphonate (GMPCPP), a slowly hydrolyzable GTP analog. After 45 min of polymerization, 1 µM taxol was added, and the reaction mixture was further incubated at 37 °C for 15 min in a water bath to stabilize the MTs. Subsequently, 1 µM tau was added, and the mixture was incubated at room temperature for a minimum of 15 minutes to allow for MT bundling.

A flow cell was prepared by adhering two glass coverslips (24 mm × 50 mm and 18 mm × 18 mm) together using two strips of double-sided tape to create a channel. The flow cell was coated by introducing 10 µL of 0.1 mg/mL bovine serum albumin (BSA) solution in BRB80 buffer, followed by a 5-minute incubation to prevent nonspecific adhesion. Afterward, MTs (either with or without tau) were introduced into the flow cell, and the samples were visualized under a TIRF microscope.

Actin filaments were polymerized by incubating 5.5 µM actin in polymerization buffer (50 mM KCl, 2 mM MgCl₂, 0.2 mM CaCl₂, and 4.2 mM ATP in 15 mM Tris-HCl, pH 8.0) at room temperature for 60 min. To prepare actin bundles, 1.2 µM fascin or α-actinin was incubated with preformed actin filaments for 10 minutes at room temperature.

A flow cell was prepared as described above and coated by introducing 10 µL of 0.1 mg/mL BSA solution in G-buffer (5 mM Tris-HCl, pH 8.0, and 0.2 mM CaCl₂), followed by a 5-minute incubation. Fascin-bundled or α-actinin-bundled actin filaments (10 µL) were then introduced into the flow cell and incubated for 5 minutes. Prior to imaging, the flow channel was washed with G-buffer to remove unbound filaments.

For experiments involving both MTs and actin filaments, a mixture of 10 µM tubulin (80% unlabeled and 20% ATTO 647-labeled) and 5.5 µM actin (90% unlabeled and 10% ATTO 488-labeled) was polymerized at 37 °C for 60 minutes. Polymerization was carried out in a buffer containing BRB80, 1 mM GMPCPP, 50 mM KCl, 2 mM MgCl₂, 0.2 mM CaCl₂, and 4.2 mM ATP, with or without the addition of tau and fascin (or α-actinin). In experiments involving the crowding agent, 0.2% (v/v) methylcellulose was included in the polymerization buffer.

### MT-actin-tau co-pelleting assay and SDS-PAGE analysis

Preformed actin filaments were incubated with preassembled MTs in BRB80 buffer for 30 min at room temperature, either in the presence or absence of 3 µM tau. Following incubation, 20 µL of each sample was transferred to an ultracentrifuge tube (Beckman Coulter) and centrifuged at ∼100,000 × *g* for 15 minutes at room temperature using an Airfuge (Beckman Coulter).

After centrifugation, 20 µL of the supernatant was carefully recovered to avoid disturbing the pellet. Both the supernatant (S) and pellet (P) fractions were mixed with 4× Laemmli loading buffer (Bio-Rad) containing 10% 2-mercaptoethanol. The samples were then heated at 90 °C for 10 min before being loaded onto a 4–20% Bis-Tris polyacrylamide gel (GenScript) for SDS-PAGE analysis. The gel was subsequently stained using SimplyBlue stain (Invitrogen) and imaged with a Sapphire Biomolecular Imager (Azure Biosystems) using excitation/emission wavelengths of 658/710 nm.

### GUV preparation

Giant unilamellar vesicles (GUVs) were generated using a modified continuous droplet interface crossing encapsulation (cDICE) technique based on an emulsion-based method^32,79^. Three distinct solutions were prepared: an inner encapsulation solution, a lipid-in-oil dispersion, and an outer aqueous solution.

For the lipid-in-oil dispersion, a 0.4 mM lipid mixture was prepared containing 49.9% 1,2-dioleoyl-sn-glycero-3-phosphocholine (DOPC), 50% poly(ethylene oxide)-poly(butadiene) (PEO-PBD), and 0.1% 1,2-dioleoyl-sn-glycero-3-phosphoethanolamine-N-(7-nitro-2-1,3-benzoxadiazol-4-yl) (NBD-PE) in a 4:1 mixture of silicone oil and mineral oil (both from Sigma-Aldrich). This lipid/oil mixture was prepared one day prior to the experiment and stored at 4 °C until use. All lipids were purchased from Avanti Polar Lipids.

Next, a reaction mixture containing 10 µM tubulin (including 20% ATTO 488-labeled tubulin) and/or 5.5 µM actin (including 10% ATTO 647-labeled actin) in polymerization buffer supplemented with GMPCPP was prepared just before encapsulation and kept on ice. Actin crosslinking proteins, either fascin or α-actinin, and tau were then added to the reaction mixture, followed by the addition of 7.5% OptiPrep density gradient medium (Sigma-Aldrich). GUVs were subsequently generated using the cDICE method.

A cDICE chamber was 3D-printed with Clear V4 resin (Formlabs) using a Form 3 3D printer (Formlabs) and mounted on the rotor of a benchtop stirrer hot plate. The chamber was rotated at 1,200 rpm. Approximately 0.7 mL of the outer aqueous solution (280 mM D-glucose, osmotically matched to the inner solution) and ∼5 mL of the lipid-in-oil dispersion were sequentially introduced into the rotating chamber. To generate GUVs, a water-in-oil emulsion was first formed by vigorously pipetting 20 µL of the reaction mixture into 0.7 mL of the lipid-in-oil dispersion. The resulting emulsion was then transferred into the rotating chamber. After 30 seconds of rotation following the addition of the emulsion droplets, the chamber was unmounted, and the GUVs were collected by recovering the aqueous outer solution into an Eppendorf tube.

Actin- or tubulin-encapsulated GUVs were incubated at 37 °C for 60 min to polymerize MTs before imaging. No taxol was used in this case. The vesicles were visualized using fluorescence microscopy.

### Fluorescence microscopy imaging

Following polymerization and incubation, MTs and actin filaments, with or without crosslinking proteins, were placed in a coverslip imaging chamber. TIRF microscopy was performed using a Nikon TiE-Perfect Focus System (PFS) microscope equipped with an Apochromat 100× objective (NA 1.49), a scientific complementary metal-oxide semiconductor (sCMOS) camera (Flash 4.0; Hamamatsu Photonics, Japan), and a laser launch controlled by an acousto-optic tunable filter. Actin and MTs were imaged using dual-color excitation with 488 nm and 647 nm lasers (Coherent Sapphire).

Fluorescence imaging of GUVs was performed using an Olympus IX-81 inverted microscope equipped with a spinning-disk confocal system (Yokogawa CSU-X1), an oil immersion 60×/1.4 NA Plan-Apochromat objective, solid-state lasers (Solamere Technology) controlled by a National Instruments DAQmx, and an iXON3 EMCCD camera (Andor Technology). Image acquisition was controlled using MetaMorph software (Molecular Devices).

Fluorescence images of lipid membranes, MTs, and actin filaments were captured using a 561 nm laser (500 ms exposure), a 647 nm laser (500 ms exposure), and a 488 nm laser (1000 ms exposure), respectively. A Semrock quad-band bandpass filter was used for emission detection.

### Image analyses

All GUV images were analyzed using Fiji (ImageJ)^80^. Fluorescence intensity profiles, presented in Figure 3b, were obtained by measuring the fluorescence intensity along the designated blue lines indicated in Figure 3a. Background signal was measured and subtracted from the intensity values, followed by normalization of the signal intensity to the maximum detected value.

For quantification of MT and actin colocalization, as presented in Figures 3d, **4b**, and **4d**, Pearson’s correlation coefficient was determined for individual GUVs using the Coloc2 plugin in Fiji (ImageJ)^80^. We used individual z-slices rather than maximum-intensity projections of z-stacks to compute Pearson’s correlation coefficient. To facilitate intuitive interpretation of the quantitative colocalization analysis, representative field-of-view images corresponding to distinct Pearson correlation coefficients are presented in **Figure S28**, highlighting how variations in coefficient values reflect differing degrees of MT-actin spatial overlap.

We analyzed the distribution of Pearson’s correlation coefficients across three independent experimental days to assess reproducibility and data consistency, revealing comparable trends and statistically consistent results under identical experimental conditions (**Figure S29**).

To analyze MT organization in GUVs, we developed a segmentation and classification pipeline (described in detail in the Supplementary Information). GUVs were identified in fluorescence images using circular detection algorithms^81,82^ combined with the CellPose segmentation algorithm^83^ and only those encapsulating MTs were retained for analysis. Local image patches centered on each vesicle were extracted from the MT channel, with external regions masked to isolate intravesicular structures. A subset of these patches was manually annotated with five exclusive structural labels: patch, filament, cluster, network, and null. A pre-trained EfficientNet-B0model^84^ was fine-tuned on the annotated dataset using focal loss^85^ and standard data augmentation. The trained classifier was then applied to the full dataset to assign structural labels, enabling quantitative comparisons across experimental conditions.

### Statistical analyses

The data presented in Figure 3B are normalized, and background subtraction is applied to remove residual background intensity prior to graph generation. All values are reported as the mean ± standard error of the mean (SEM). Each experiment was conducted with a minimum of three independent replicates.

Statistical analyses and graph generation were performed using GraphPad Prism. Depending on the dataset, statistical significance was assessed using an unpaired t-test or one-way and two-way ANOVA, followed by Tukey’s Honest Significant Difference post-hoc test for multiple comparisons. The calculated *p*-values are provided in Table S1 in the Supporting Information.

### Calculation of persistence length

The persistence length of MT and actin bundles was determined by measuring the end-to-end distance and contour length of individual filaments by ImageJ using the Analyze-Measure option^71,72^. The data were analyzed by fitting to the following equation:

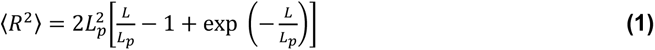

where 〈*R*^2^〉 is the mean squared end-to-end distance, *L* is the contour length of the bundle, and L_p_, represents the persistence length of the bundle. The derived parameter from the fitting gave the values of persistence lengths (**Figure S5**).

### Simulations

Langevin dynamics simulations were performed using the open-source molecular dynamics simulator LAMMPS. Systems contained two species of semiflexible filaments with bending rigidities κ_stiff_ and κ_soft_, representing MTs and actin bundles, respectively. Filaments were modeled as bead-spring chains, with harmonic stretching potentials between adjacent beads and harmonic angle potentials acting on adjacent bead triplets. Membranes were modeled using the particle-based coarse-grained model described in ref.^60^. Nonbonded interactions—both filament-filament and filament-membrane were modeled using purely repulsive Weeks-Chandler-Andersen potentials^86^. “Sticky” crosslinker sites were uniformly distributed along polymer chains with a specified crosslinker coverage fraction ϕ and were permitted to form reversible harmonic bonds with compatible sites through a probabilistic Monte Carlo scheme. Systems were equilibrated prior to activation of crosslinking. The crosslinker coverage fraction ϕ and filament bending rigidities were systematically varied to explore parameter space. Complete parameter sets and implementation details are provided in Supplementary Information.

## RESOURCE AVAILABILITY

### Lead contact

Requests for further information and resources should be directed to and will be fulfilled by the lead contact, Aaron Dinner and Allen Liu.

### Materials availability

Details on commercially available materials are given in the text.

### Data and code availability

Raw image data will be made available by request. The code for microtubule structure classification and analysis is available at: https://github.com/Center-for-Living-Systems/structure_classification. Code for the simulations is available at https://github.com/jordanshivers/cytoskeletal-network-assembly.

## ACKNOWLEDGMENTS

We acknowledge support from the National Science Foundation to APL (MCB 2201236) and ARD (MCB 2201235) and from the National Institutes of Health to APL (NIBIB R01EB030031). ARD and LD acknowledge support from the Physics Frontier Center for Living Systems (CLS) funded by the National Science Foundation (PHY-2317138). JLS acknowledges support from the Eric and Wendy Schmidt AI in Science Postdoctoral Fellowship, a Schmidt Sciences program. We thank D. Kovar (University of Chicago) for α-actinin.

## AUTHOR CONTRIBUTIONS

MA and APL designed the experiments. MA conducted the experiments and analyzed the experimental data. JS and ARD designed the modeling and simulations. JLS conducted the simulations and analyzed the simulation data. LD developed the image-analysis methods and analyzed the MT architectures in GUV. MA, JS, ARD, and APL wrote the initial draft of the paper. All authors discussed the results and edited the paper.

## DECLARATION OF GENERATIVE AI AND AI-ASSISTED TECHNOLOGIES

The authors declare no use of AI and AI-assisted technologies.

## SUPPLEMENTAL INFORMATION

**Document S1. Supplemental methods, Figures S1–S29, and Tables S1–S2**

