## Supplementary Information for "Tau-Driven Coordination of Microtubule-Actin Crosstalk in Cell-Sized Vesicles"

#### 1. Probability distribution of different microtubule architectures in GUVs

##### (a) GUV-encapsulated microtubule structure segmentation

Microtubule (MT) channel images were first preprocessed to enhance signal quality and reduce artifacts. A median filter was applied to remove bright spot noise, followed by background subtraction using the median image to correct for uneven illumination. The images were then normalized to balance dynamic range. To identify GUVs that encapsulate MTs, the GUV channel was analyzed using a multi-scale Hough transform to detect circular objects. Detected circles were screened to exclude incomplete objects touching image boundaries and those with sizes outside a defined range. Additionally, each candidate region was evaluated in the MT channel by computing the standard deviation of pixel intensities; regions with low standard deviation, indicating insufficient MT content, were excluded. For each remaining object, a local image patch was cropped from the MT channel, and the region outside the GUV segmentation was filled with the background intensity level of the MT channel. This masking step effectively removed signal contributions from neighboring structures and ensured that downstream analysis focused solely on the encapsulated content. The final set of segmented circular regions represents GUV-encapsulated MTs used for subsequent structural classification and statistical analysis. This workflow is illustrated with an example in **Figure S2**.

##### (b) MT structure classification and analysis

A dataset comprising 1,064 image patches (each 96 × 96 pixels) was extracted from microscopy experiments conducted across multiple days and tau concentration conditions. From this collection, a randomly selected subset of 335 patches was manually annotated into five structural classes: patch, filament, cluster, network, and null. The first four classes correspond to distinct MT organizations, while the null class represents patches lacking sufficient MT content.

For classification, a pre-trained EfficientNet-B0 model[1] was fine-tuned on the annotated dataset. Input images, originally grayscale, were converted to three-channel RGB and resized to 224 × 224 pixels. Standard data augmentations, including random horizontal flips, rotations (±10°), and color jitter, were applied to enhance generalization. The model's classifier head was modified to include a dropout layer ($p = 0.3$) followed by a fully connected layer matching the number of classes. To mitigate class imbalance and emphasize difficult examples, we employed a focal loss function ($\alpha = 1$, $\gamma = 2$). Optimization was performed using the AdamW optimizer[2] (learning rate = $3\times10^{-4}$, weight decay = $1\times10^{-4}$) combined with a cosine annealing learning rate scheduler. The dataset was split into training (80%) and validation (20%) sets using stratified sampling, and additional stratified oversampling was applied to balance class representation during training. Training proceeded with early stopping based on validation accuracy, and the model achieving the highest validation performance was retained for evaluation. Classification results are presented in **Figure S3**. The trained classifier was applied to the full dataset to assign predicted class labels to all image samples.

### 2. Simulation details

Simulation units are specified in terms of characteristic scales of energy $\varepsilon$, mass $m$, and distance $\sigma$. Particles of all types are assigned mass $m$. Simulations are performed using the open-source molecular dynamics simulator LAMMPS[3], using a Langevin thermostat with unit reduced temperature and unit damping. The simulation timestep is $\Delta t = 0.0025\tau_0$, in which $\tau_0 = \sigma\sqrt{m/\varepsilon}$. Parameter values are specified in Table S2.

#### (a) Membrane model

To model flexible fluid vesicles, we use a particle-based coarse-grained membrane model[4]. This model represents a bilayer membrane with oriented spherical particles interacting via a pairwise Lennard-Jones-like potential that favors the alignment of neighboring particles. This model has been used extensively in prior work[5-9]. We adapt the LAMMPS implementation of this model developed by Forster *et al.*[9]

Each spherical membrane particle, with diameter $\sigma_m = \sigma$, is characterized by a position vector $\vec{r}_i$ and unit orientation vector $\hat{n}_i$, as illustrated schematically in **Figure S10**. Pairs of membrane particles $i$ and $j$ interact via an anisotropic pair potential $U_{\mathrm{mem}}(\vec{r}_{ij}, \hat{n}_i, \hat{n}_j)$ defined as

$$U_{\mathrm{mem}}(\vec{r}_{ij}, \hat{n}_i, \hat{n}_j) = \begin{cases} u_{\mathrm{R}}(r) + [1 - \phi(\hat{r}_{ij}, \hat{n}_i, \hat{n}_j)]\varepsilon_{\mathrm{m}}, & r < r_{\mathrm{min}} \\ u_{\mathrm{A}}(r)\phi(\hat{r}_{ij}, \hat{n}_i, \hat{n}_j), & r_{\mathrm{min}} < r < r_c^{\mathrm{m}} \end{cases}$$

in which $\vec{r}_{ij} = \vec{r}_j - \vec{r}_i$, $r = |\vec{r}_{ij}|$, and $\hat{r}_{ij} = \vec{r}_{ij}/r$, and the repulsive and attractive components are defined as

$$u_{\mathrm{R}}(r) = \varepsilon_{\mathrm{m}} \left[ \left(\frac{r_{\mathrm{min}}}{r}\right)^4 - 2\left(\frac{r_{\mathrm{min}}}{r}\right)^2 \right]$$

and

$$u_{\mathrm{A}}(r) = -\varepsilon_{\mathrm{m}} \left( \cos\left[ \frac{\pi}{2} \frac{(r - r_{\mathrm{min}})}{(r_c^{\mathrm{m}} - r_{\mathrm{min}})} \right] \right)^{2\zeta}.$$

As in Yuan *et al*[2]. the distance corresponding to the potential minimum is set to $r_{\mathrm{min}} = 2^{1/6}\sigma_{\mathrm{m}}$ and the cutoff distance is set to $r_c^{\mathrm{m}} = 2.6\sigma_{\mathrm{m}}$, such that next-nearest-neighbor interactions are considered. The exponent $\zeta$ determines the slope of the attractive branch, thereby influencing the in-plane diffusivity of membrane particles. A higher $\zeta$ value corresponds to an increased diffusivity. The attractive branch decays smoothly to zero at $r = r_c^{\mathrm{m}}$. The dimensionless function $\phi(\hat{r}_{ij}, \hat{n}_i, \hat{n}_j)$ introduces an aligning tendency between neighboring membrane particles and is defined as

$$\phi(\hat{r}_{ij}, \hat{n}_i, \hat{n}_j) = 1 + \mu\left[\mathrm{a}(\hat{r}_{ij}, \hat{n}_i, \hat{n}_j) - 1\right]$$

in which the parameter $\mu$ is a dimensionless aligning stiffness that governs the emergent bending rigidity $\kappa_{\mathrm{m}}$ of the resulting membrane, and the function $\mathrm{a}(\hat{r}_{ij}, \hat{n}_i, \hat{n}_j)$ is defined as

$$\mathrm{a}(\hat{r}_{ij}, \hat{n}_i, \hat{n}_j) = (\hat{n}_i \times \hat{r}_{ij}) \cdot (\hat{n}_j \times \hat{r}_{ij}) + \sin\theta_0\, (\hat{n}_j - \hat{n}_i) \cdot \hat{r}_{ij} - \sin^2\theta_0,$$

in which $\theta_0$ corresponds to the preferred spontaneous curvature angle. We consider the case of zero spontaneous curvature ($\theta_0 = 0$) and choose a value of $\mu$ that yields a macroscopic membrane bending rigidity of $\kappa_{\mathrm{m}} \approx 18k_{\mathrm{B}}T$.

For all simulations involving membrane particles, we prepare and equilibrate a vesicle of average radius $R$ within a periodic box of side length $L$ prior to the placement of internal polymers.

**(b) Polymer model**

We next introduce semiflexible polymers composed of spherical particles of diameter $\sigma_p$ connected by Hookean springs of rest length $l_0$ with spring constant $k_{\mathrm{stretch}}^{\alpha}$, with triplets of adjacent particles interacting via a harmonic bending interaction with spring constant $k_{\mathrm{bend}}^{\alpha}$, in which $\alpha \in \{\mathrm{soft}, \mathrm{stiff}\}$ corresponds to one of the two polymer species. The polymer stretching energy for species $\alpha$ is given by

$$U_{\mathrm{stretch}}^{\alpha} = \sum_{ij} k_{\mathrm{stretch}}^{\alpha} (l_{ij} - l_0)^2,$$

in which the sum is taken over all pairs of adjacent particles $i$ and $j$ along the polymer backbone. The bending energy for species $\alpha$ is given by

$$U_{\text{bend}}^{\alpha} = \sum_{ijk} k_{\text{bend}}^{\alpha} \left(\theta_{ijk} - \theta_0^{\text{p}}\right)^2,$$

in which the sum is taken over all triplets of adjacent particles $i$, $j$, and $k$ and the rest angle is given by $\theta_0^{\text{p}} = 0$. Note that the bending spring constant for species $\alpha$ is proportional to the polymer bending rigidity $\kappa_\alpha = 2l_0 k_{\text{bend}}^{\alpha}$, and persistence length $l_p^{\alpha} = \kappa_\alpha / k_B T$. All polymer particles interact via a pairwise purely repulsive Weeks-Chandler-Andersen (WCA) potential[10], defined as

$$U_{\text{WCA}}^{\text{p}}(r) = \begin{cases} 4\varepsilon_{\text{p}} \left[ \left(\frac{\sigma_{\text{p}}}{r}\right)^{12} - \left(\frac{\sigma_{\text{p}}}{r}\right)^{6} \right] + \varepsilon_{\text{p}}, & r \leq 2^{1/6}\sigma_{\text{p}} \\ 0, & r > 2^{1/6}\sigma_{\text{p}} \end{cases}.$$

Likewise, a pairwise purely repulsive WCA interaction acts between membrane particles and polymer particles, with

$$U_{\text{WCA}}^{\text{mp}}(r) = \begin{cases} 4\varepsilon_{\text{mp}} \left[ \left(\frac{\sigma_{\text{mp}}}{r}\right)^{12} - \left(\frac{\sigma_{\text{mp}}}{r}\right)^{6} \right] + \varepsilon_{\text{mp}}, & r \leq 2^{1/6}\sigma_{\text{mp}} \\ 0, & r > 2^{1/6}\sigma_{\text{mp}} \end{cases},$$

in which $\sigma_{\text{mp}} = \left(\sigma_{\text{p}} + \sigma_{\text{m}}\right)/2$.

The lengths of the soft and stiff filaments are specified as $L_\alpha = (n_\alpha - 1)l_0$, in which $n_\alpha$ is the number of particles per filament of species $\alpha$. The total number of particles of species $\alpha$ is given by $N_\alpha = \rho_\alpha V$, in which $\rho_\alpha$ is the number density of particles of species $\alpha$ and $V$ is the volume of the space into which the particles are to be placed (for bulk simulations, $V = {L_{\text{box}}}^3$, in which $L_{\text{box}}$ is the side length of the cubic periodic simulation box, whereas for vesicle-confined simulations, $V \approx \frac{4}{3}\pi R^3$).

Filaments are initially placed with random positions and orientations within the specified target volume. Then, a brief simulation is carried out in which the WCA polymer-polymer interaction potential is replaced with a soft repulsive potential to remove particle overlaps. Next, the WCA polymer-polymer interaction potential is reinstated, and the system is equilibrated for $\tau_{\text{equil}} = 2 \times 10^5 \Delta t$. Finally, the formation of crosslinks is enabled and the main simulation stage proceeds for duration $\tau_{\text{main}} = 1.5 \times 10^6 \Delta t$.

#### 3. Crosslinking procedure

To model crosslinking, we adapt a similar approach to Ref. [3], in which a fraction of randomly chosen polymer particles is marked as linker-decorated, i.e., “sticky” particles capable of forming a crosslink with another particle. Specifically, during initialization, we randomly select a fraction $\phi_\alpha$ of particles of each species α to be sticky sites. Site designations are specified during initialization and remain fixed throughout the simulation. During each timestep, for any pair of polymer particles of species $\alpha$ and $\beta$ in which (1) one or both particles is sticky and (2) the inter-particle distance is less than $r_c^{\mathrm{p}}$, a new crosslink is added with probability $p_{\mathrm{cl}}^{\alpha,\beta}$. At the end of each timestep, crosslinks are randomly broken with probability $p_{\mathrm{cl}}^{\mathrm{break}}$. Each crosslink is modeled as a harmonic spring with rest length $l_0^{\mathrm{cl}}$ and spring constant $k_{\mathrm{stretch}}^{\mathrm{cl}}$. Each particle is allowed to form a maximum of $\nu_{\max} = 3$ crosslinks.

For a given filament, each sticky site represents a segment of the filament with bound crosslinker molecules. As discussed in Syed *et al*,[11] our objective is to represent an idealized situation in which all crosslinker molecules have become bound to filaments and none remain in solution. In this scenario, the specified coverage fraction ϕ of sticky sites acts as a surrogate for the experimentally controlled concentration of crosslinker (tau) molecules.

#### 4. Visualization of simulated configurations

Simulation configurations are rendered using *Ovito*[12]. To highlight network architecture and enhance crosslink visibility, spherical polymer particles are concealed from view in most figures. Only the corresponding polymer bonds (magenta for the stiff polymer species and green for the soft polymer species) and crosslink bonds (black) are displayed. **Figure S11** displays an example of a bulk network configuration with and without visible particles.

To aid in the visualization of the internal network structure in simulations of vesicle-confined networks, we replace the membrane particles with a surface mesh computed from the particle positions and then “cut away” half of the mesh closest to the viewer. **Figure S12** shows an example of a vesicle configuration with visible membrane particles, as well as the corresponding configuration in which the membrane particles are replaced by a surface mesh.

### 5. Colocalization analysis of simulation configurations

#### (a) Colocalization method

To quantify the dependence of the colocalization of the two polymer species on simulation parameters, we generate artificial fluorescence images representing different simulation configurations. We then perform colocalization analysis on these images, using the same technique employed for the experimental images.

We first divide the simulation domain into $d_x \times d_y \times d_z$ voxels. The fluorescence intensity contributed to a given voxel by a nearby particle corresponds to the value, at the voxel midpoint, of a three-dimensional Gaussian distribution of unit amplitude and standard deviation $\sigma = 0.5$ centered on the particle position $(r_{i,x}, r_{i,y}, r_{i,z})$. In **Figure S13**, we show a $z$-slice of the intensity distribution corresponding to a representative simulated periodic network configuration, with stiff polymers shown in magenta and soft polymers in green, as well as an overlay of both channels. We also show the corresponding sum and maximum intensity projections, which correspond to the total pixel intensity and maximum pixel intensity, respectively, calculated along the $z$-axis. In **Figure S14**, we show a corresponding set of artificial fluorescence images for a representative simulated vesicle-confined network, in which the fluorescence intensity of membrane particles is shown in red.

To measure the colocalization of the two polymer species, we compute the Pearson's correlation coefficient between the pixel intensities for stiff and soft polymer particles across all $z$-slices. We have verified that identical values of the Pearson's correlation coefficient are obtained when the same simulation-derived images are analyzed using the Coloc2 plugin in Fiji (ImageJ)[13].

#### (b) Influence of confinement, crosslinker concentration, and filament properties on colocalization

We find that, when crosslinkers are present, increasing the bending stiffness of the soft polymer species increases colocalization between the soft and stiff polymer species (see **Figure 5** in the main text). This observation supports our argument that when both species have sufficiently large persistence lengths, the formation of composite bundles consisting of aligned polymers of both species becomes favorable. **Figure S15** shows a close-up view of a bundle of aligned stiff and soft polymers observed in a system where this condition is met, with $\kappa_{\text{soft}}/\kappa_{\text{stiff}} = 0.5$.

**Figure S16** shows the final configurations of bulk systems across a wide range of values of the crosslinker coverage fraction $\phi$ and bending stiffness ratio $\kappa_{\text{soft}}/\kappa_{\text{stiff}}$. When $\kappa_{\text{soft}}/\kappa_{\text{stiff}}$ is small, bundles of aligned stiff polymers are still consistently observed, but bundles containing a mixture of aligned stiff and soft polymers are significantly less likely to form.

#### 6. Exploration of parameter space

The "phase space portraits" in **Figures S17-24** show the influence of various parameters on the structural characteristics of the final network assemblies ($\tau_{\text{main}} = 1.5 \times 10^6 \Delta t$). The values of any parameters not explicitly specified here are given in **Table S2**.

**Figures S16** and **S17** show bulk configurations for system sizes $L_{\text{box}} = 40\sigma$ and $L_{\text{box}} = 60\sigma$, respectively, with varying crosslinker coverage fraction $\phi$ and bending stiffness ratio $\kappa_{\text{soft}}/\kappa_{\text{stiff}}$, for species concentrations $\rho_{\text{soft}} = \rho_{\text{stiff}} = 0.05\sigma^{-3}$ . Systems with $L_{\text{box}} = 60\sigma$ and lower species concentrations $\rho_{\text{soft}} = \rho_{\text{stiff}} = 0.025\sigma^{-3}$ and $\rho_{\text{soft}} = \rho_{\text{stiff}} = 0.0125\sigma^{-3}$ are shown in **Figure S18** and **S19**, respectively.

**Figure S20** shows examples of networks assembled in vesicles of radius $R = 27\sigma$ with polymer concentrations $\rho_{\text{soft}} = \rho_{\text{stiff}} = 0.05\sigma^{-3}$, in which for large $\phi$ and large $\kappa_{\text{soft}}/\kappa_{\text{stiff}}$, the formation of aligned bundles containing both polymer species is clearly visible. Although their occurrence is less clearly pronounced, bundles containing both species can also be seen for some values of $\phi$ and $\kappa_{\text{soft}}/\kappa_{\text{stiff}}$. As is shown in **Figure S21**, the formation of aligned bundles containing both species is more readily observed in systems in which the polymer concentration is lowered to $\rho_{\text{soft}} = \rho_{\text{stiff}} = 0.025\sigma^{-3}$.

**Figure S22** depicts vesicle-confined assemblies with varying vesicle radius $R$ and crosslinker coverage fraction $\phi$. Here, the polymer concentrations are $\rho_{\text{soft}} = \rho_{\text{stiff}} = 0.05\sigma^{-3}$ and the bending stiffness ratio is $\kappa_{\text{soft}}/\kappa_{\text{stiff}} = 0.5$.

**Figures S23** and **S24** depict, respectively, bulk and vesicle-confined assemblies with varying filament length $L_\alpha$ and crosslinker coverage fraction $\phi$. Here, the polymer concentrations are $\rho_{\text{soft}} = \rho_{\text{stiff}} = 0.05\sigma^{-3}$ and the bending stiffness ratio is $\kappa_{\text{soft}}/\kappa_{\text{stiff}} = 0.5$.

In **Figure S25**, we plot the Pearson's correlation coefficient for bulk systems with a 1:1 polymer species ratio for three different values of the polymer concentration $\rho_\alpha$. We find that the Pearson's correlation coefficient increases as the polymer concentration $\rho_\alpha$ decreases. However, the tendency of the colocalization to increase with both crosslinker coverage fraction $\phi$ and bending stiffness ratio $\kappa_{\mathrm{soft}}/\kappa_{\mathrm{stiff}}$ remains consistent.

In **Figure S26**, we plot the Pearson's correlation coefficient for the same systems as a function of filament length $L_\alpha$. For large crosslinker coverage fraction $\phi$, colocalization decreases monotonically with increasing filament length. This behavior likely reflects the length-dependent restricted diffusion of stiff filaments in the entangled, high-concentration regime[14,15]. For systems with low but nonzero crosslinker coverage fraction $\phi$, however, colocalization is maximized at an intermediate value of the filament length. The reduced colocalization in the regime below this optimum likely reflects the fact that, for small $\phi$ and small $L_\alpha$, many filaments have either no available crosslinking sites or fewer than the two sites required for crosslinking-induced alignment.

Recall that the filament length is related to the number of constituent particles $n_\alpha$ as $L_\alpha = (n_\alpha - 1)l_0$. For filaments composed of $n_\alpha$ particles, each with crosslinker coverage fraction $\phi$, the probability that a given filament has no available crosslinking sites is $P(0) = (1-\phi)^{n_a}$. Although $P(0)$ decreases rapidly with increasing $\phi$ or $n_a$, it can be substantial for small values of either parameter. For example, for $\phi = 0.2$ and $n_a = 5$, we find $P(0) \approx 0.33$, meaning that roughly 33% of filaments cannot form crosslinks. This is consistent with the visible presence of crosslink-free filaments in the corresponding representative configurations shown in **Figures S23** and **S24**. Crosslinking-induced alignment between two filaments requires at least two crosslinks. The probability of a filament having exactly one available site is $P(1) = n_a\phi(1-\phi)^{n_a-1}$, so the probability of having fewer than two sites is $P(0 \text{ or } 1) = P(0) + P(1) = (1-\phi)^{n_a} + n_a\phi(1-\phi)^{n_a-1}$. For $\phi = 0.2$ and $n_a = 5$, this yields $P(0 \text{ or } 1) \approx 0.73$, indicating that 73% of filaments are incapable of undergoing crosslinking-induced alignment. This fraction decreases rapidly with increasing $\phi$ or $n_a$.

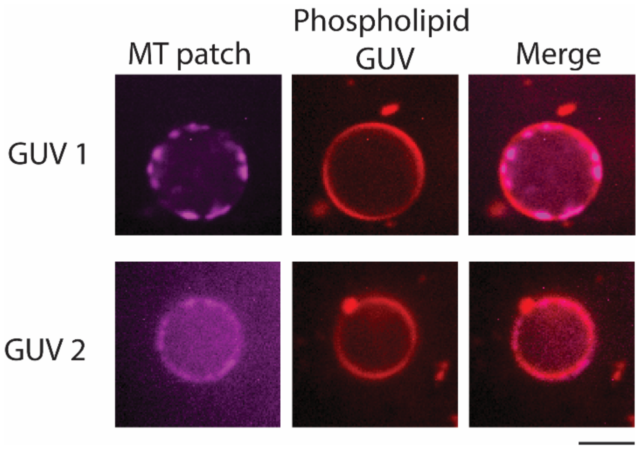


**Figure S1. Representative confocal fluorescence microscopy images showing MT patch formation within GUVs.** Scale bar: 10 μm.

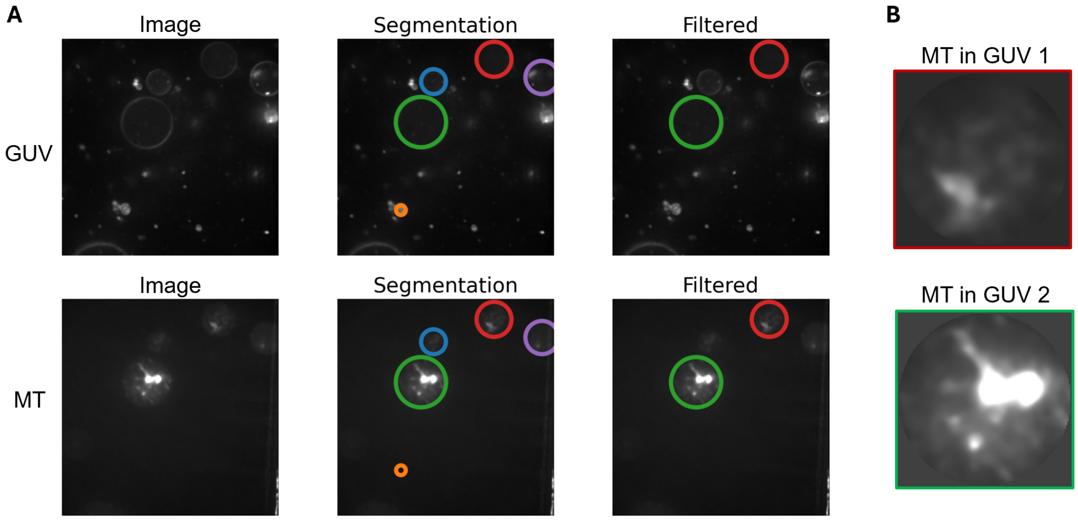


**Figure S2. Segmentation and filtering of MT-encapsulated GUV structures.** (A) The top row displays the GUV fluorescence channel, and the bottom row shows the corresponding normalized MT channel. The left column presents the original input images. The middle column overlays the initial circular object segmentations obtained from the GUV channel. The right column shows the final filtered objects after applying size, boundary, and MT content-based exclusion criteria. Colored circles indicate individual segmented regions. (B) The two MT structure image patches with regions outside the GUV filled with background intensity.

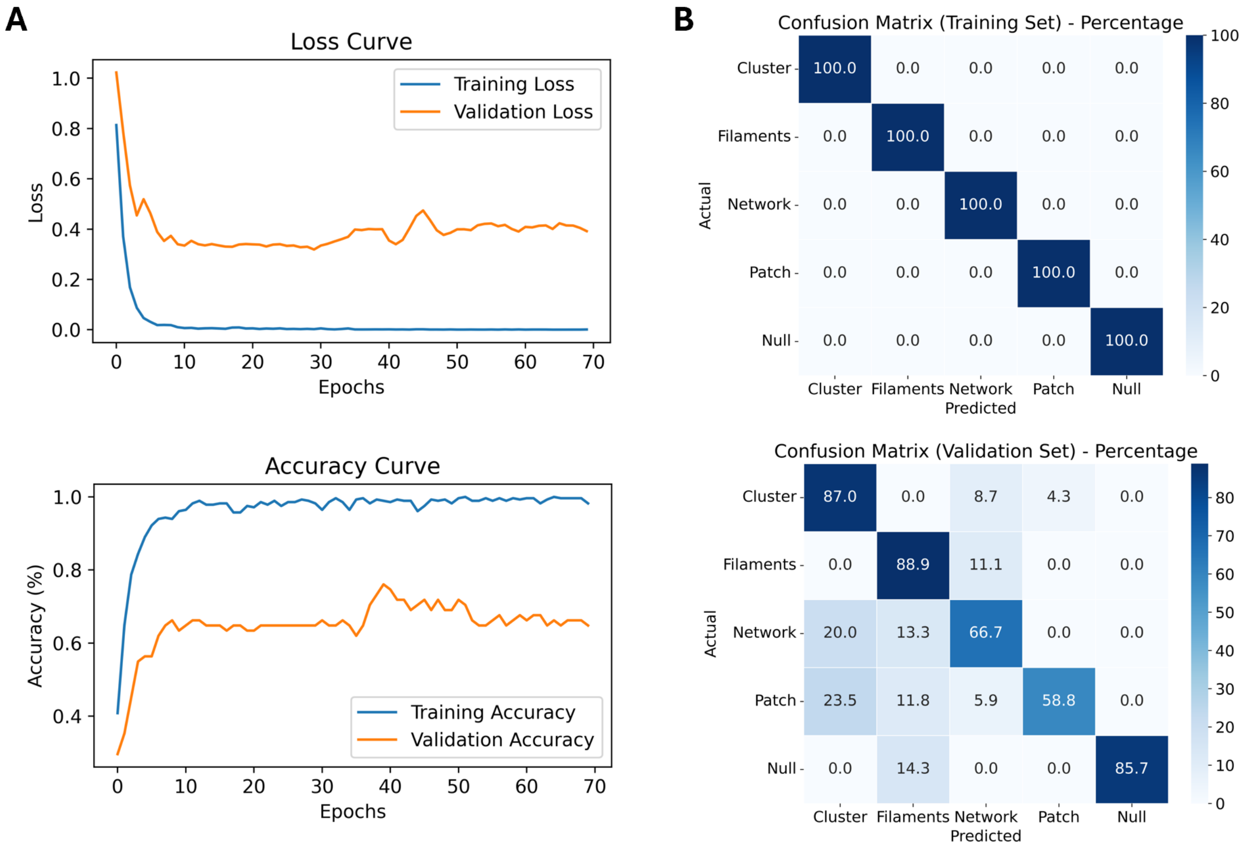


**Figure S3. Performance of the MT structure classifier.** (A) Accuracy and loss curves over training epochs for both training and validation sets. (B) Confusion matrices showing classification performance as percentages on the training set (top) and validation set (bottom).

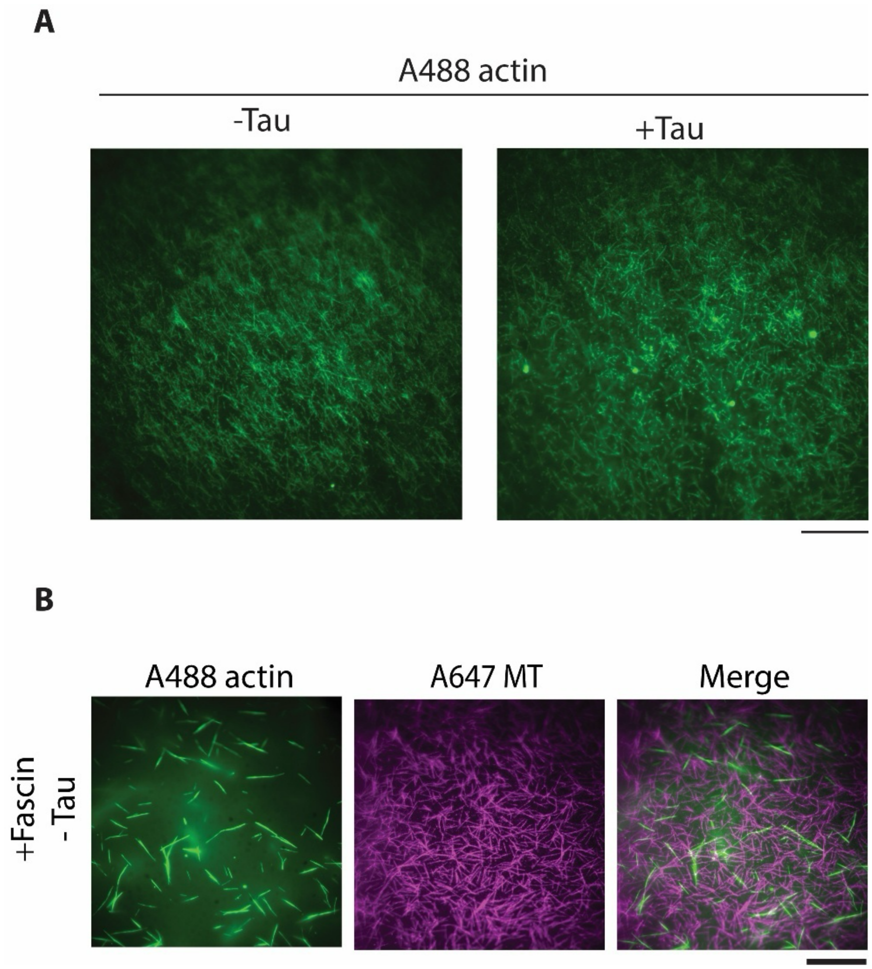


**Figure S4. Representative TIRF microscopy images depicting** (A) actin organization in the absence or presence of 2 μM tau. Actin concentration is 5.5 μM in the experiment**.** Scale bar: 20 μm. (B) MTs and fascin-mediated actin filaments in the absence of tau. Fascin concentration is 1.2 μM. Scale bar: 10 μm.

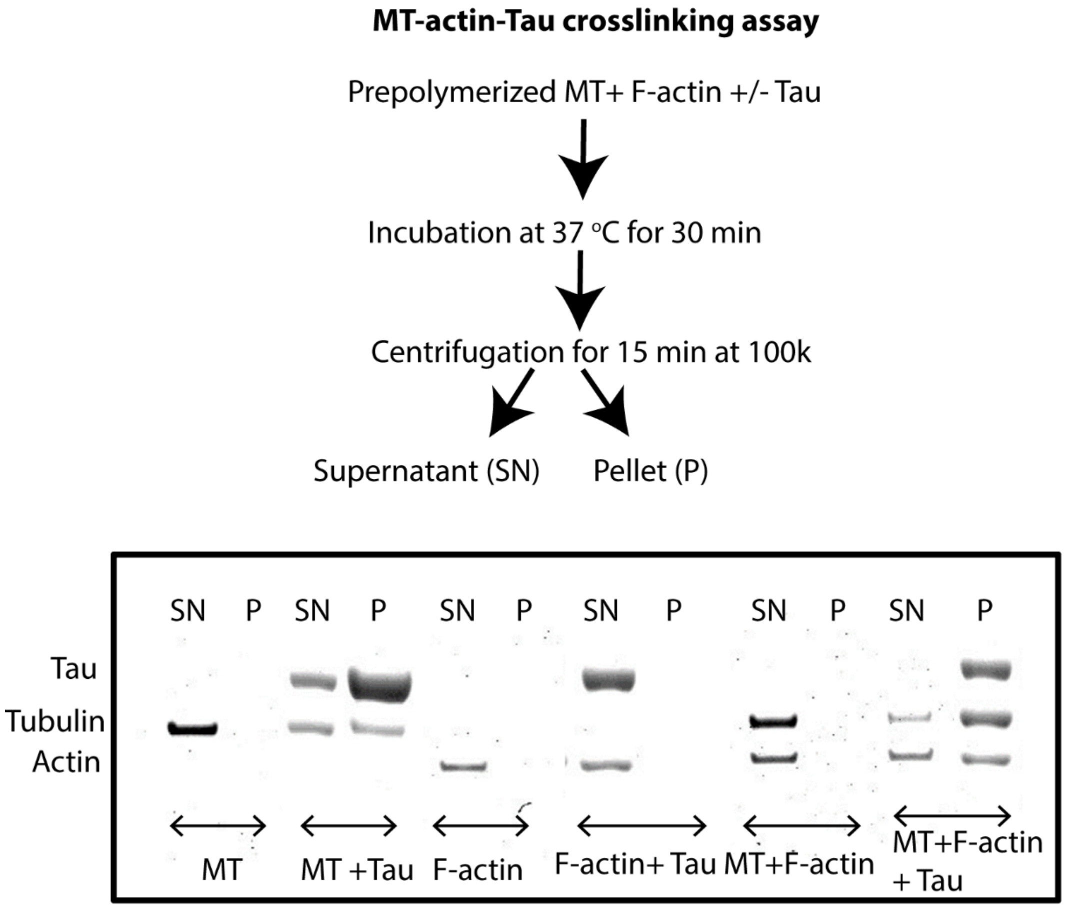


**Figure S5. Tau-mediated crosslinking of MTs and F-actin.** A crosslinking assay was performed using prepolymerized GMPCPP-stabilized MTs and F-actin, incubated with tau (2 μM). Tau was mixed with MTs (2 μM), F-actin (2 μM), or both, followed by high-speed centrifugation using an Airfuge. Supernatant (SN) and pellet (P) fractions were analyzed by SDS-PAGE. A representative SDS-PAGE gel image of pellet and supernatant fractions from the crosslinking assay is shown at the bottom of the figure. F-actin was detected in the pellet only when co-incubated with both MTs and tau, indicating tau-induced formation of MT-actin complexes.

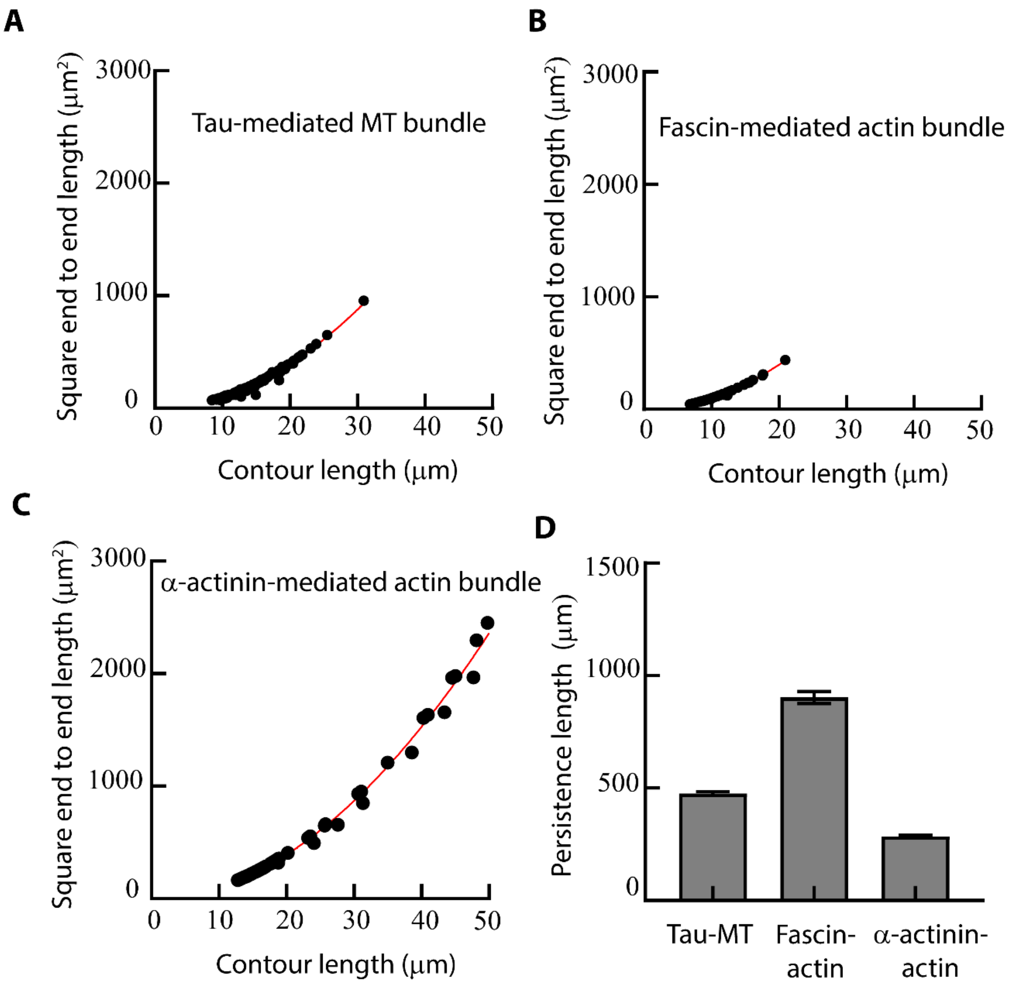


**Figure S6. Squared end-to-end lengths and contour lengths of (A) tau-mediated MT bundles, (B) fascin-mediated actin bundles, and (C) α-actinin-mediated actin bundles, respectively.** Solid red lines are fits from Equation 1 (see main text). The persistence lengths of tau-induced MT bundles, fascin-induced actin bundles, and α-actinin-induced bundles were 475.7±8.9 μm, 902.8±27.0 μm, and 285.3±6.3 μm, respectively. Error bar: standard error of the means. A minimum of 150 bundles was analyzed for each condition.

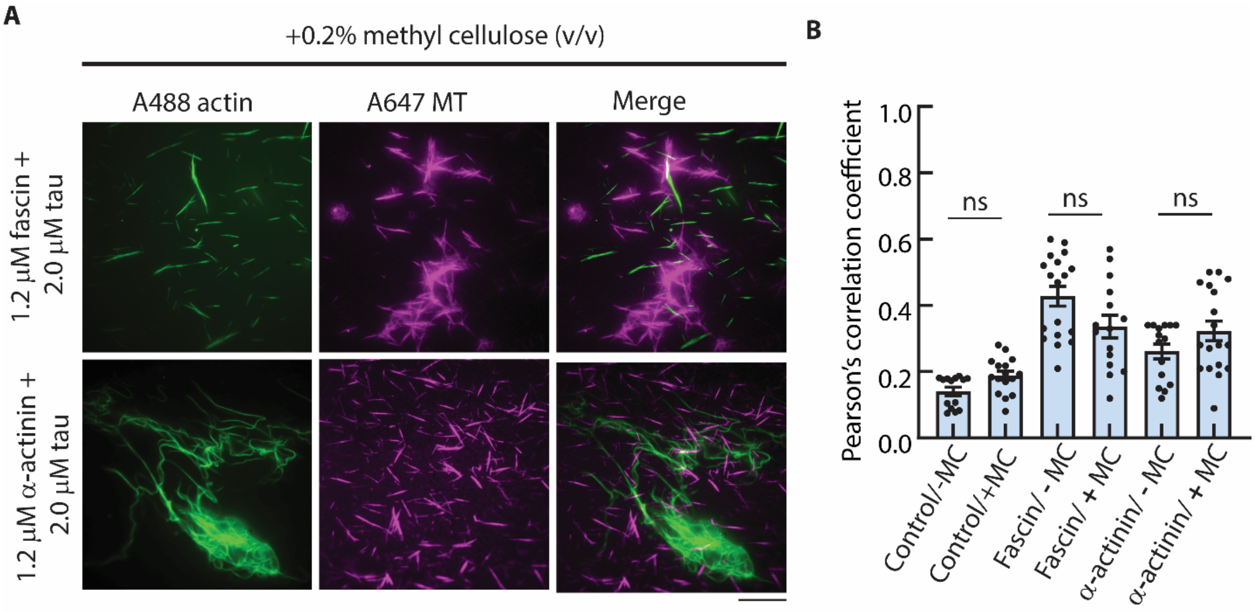


**Figure S7: MT–actin colocalization in bulk experiments performed in the presence of 0.2% (v/v) methylcellulose**. (A) Representative TIRF microscopy images showing MT–actin colocalization in the presence of 1.2 µM fascin (top) and 1.2 µM α-actinin (bottom), with 2.0 µM tau included in both conditions. Scale bar: 20 µm. (B) Comparison of Pearson's correlation coefficients of MT-actin colocalization in the presence of different associated proteins, with and without methyl cellulose. "Control" refers to only MT and actin filaments, without associated proteins. ns = non-significant difference.

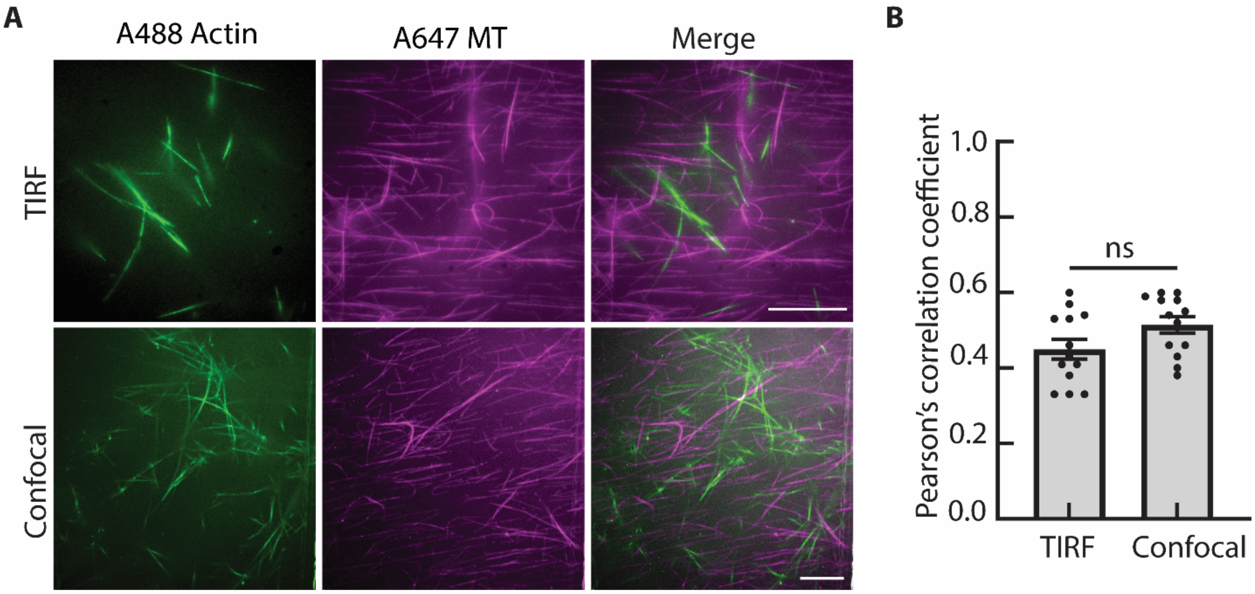


**Figure S8: Control analysis comparing MT-actin colocalization measured in bulk experiments under identical experimental conditions.** (A) Representative fluorescence microscopy images showing MT-actin colocalization acquired by TIRF and confocal microscopy at 2 μM tau and 2 μM fascin concentrations. Scale bar: 20 μm. (B) Comparison of Pearson's correlation coefficients obtained from TIRF and confocal images. ns = non-significant difference.

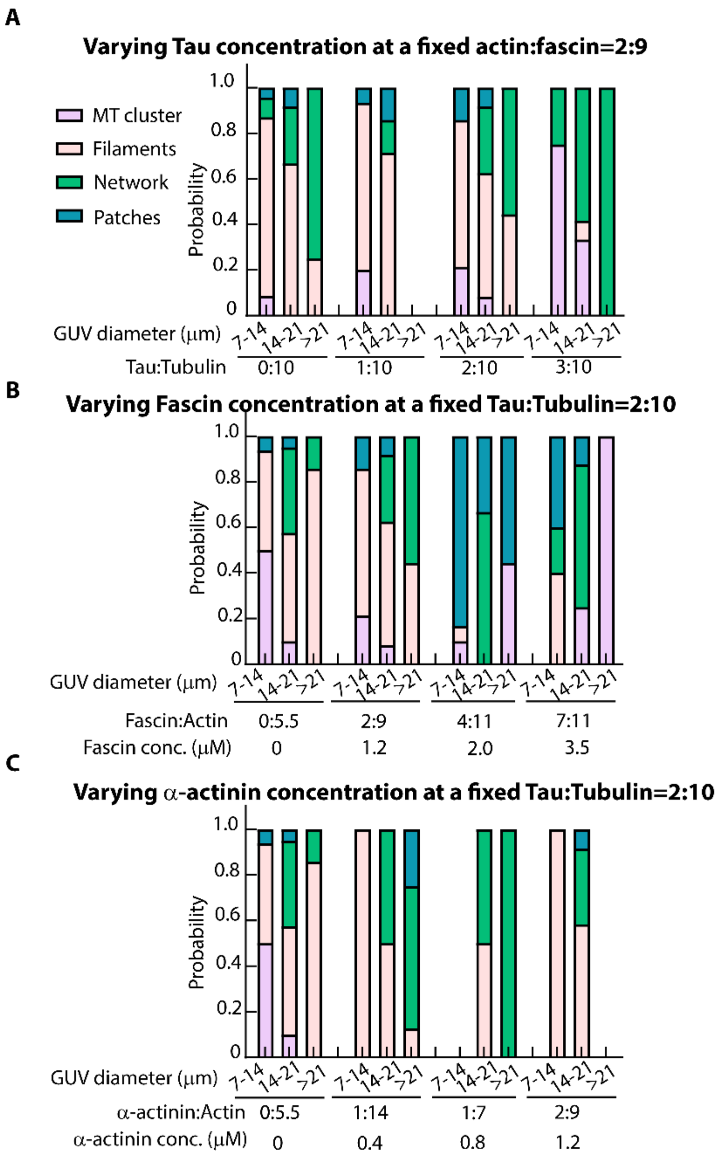


**Figure S9. Cumulative probability of different MT architectures in GUVs of varying diameters at different** (A) Tau:tubulin ratios at a fixed actin:fascin ratio of 2:9, (B) actin:fascin ratios at a fixed tau:tubulin ratio of 2:10 and (C) actin:α-actinin ratios at a fixed tau:tubulin ratio of 2:10. For analysis, the total numbers of GUVs analyzed are: A, 255; B, 146; and C, 145. Only GUVs containing both actin and MT signals were included in the analysis.

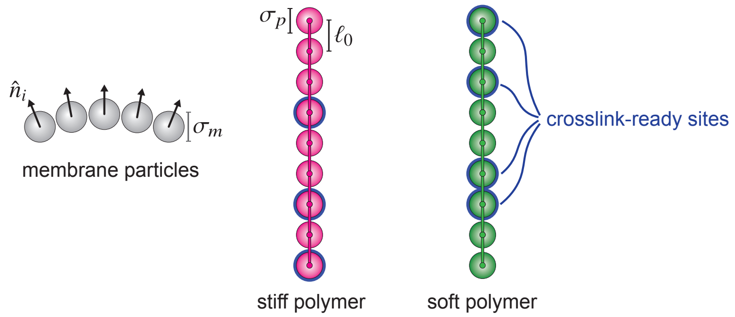


**Figure S10. Schematic of the membrane and polymer models.** (Left) Membrane particles are spheres of diameter $\sigma_m$ with associated positions $\vec{r}_i$ and orientations $\hat{n}_i$. (Center and right) Stiff (magenta) and soft (green) semiflexible polymers consisting of spherical particles with diameter $\sigma_p$ connected by harmonic springs of rest length $l_0$. Stiff and soft polymers have identical stretching constants ($k_{\text{stretch}}^{\text{stiff}} = k_{\text{stretch}}^{\text{soft}}$) but distinct bending constants ($k_{\text{bend}}^{\text{stiff}} > k_{\text{bend}}^{\text{soft}}$). Blue outlines indicate randomly assigned crosslink-ready sites with coverage fraction $\phi_\alpha$ for polymer species $\alpha$.

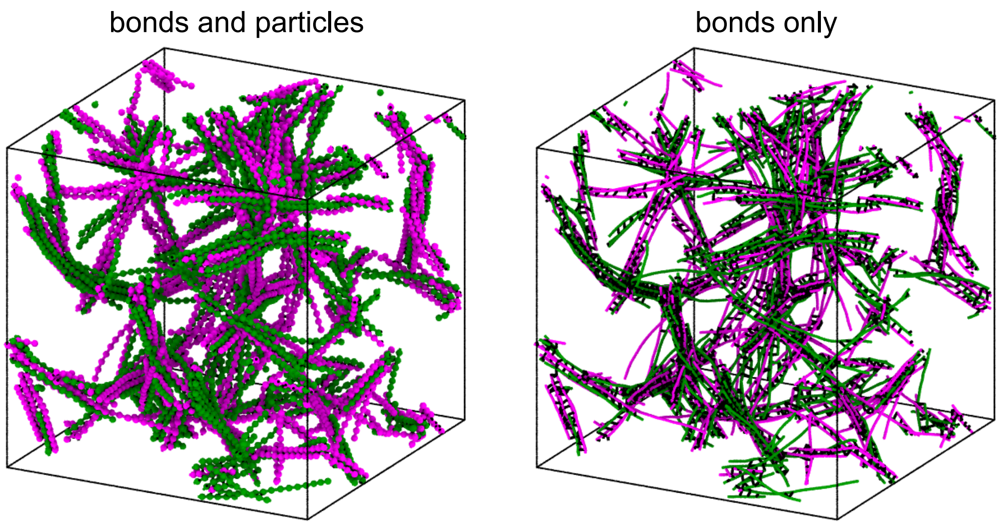


**Figure S11. Visualization of semiflexible polymer networks from simulations**. (Left) Configuration displaying both bonds and spherical polymer particles. (Right) The same configuration with bonds only, omitting particles to enhance the visibility of crosslinks (black bonds).

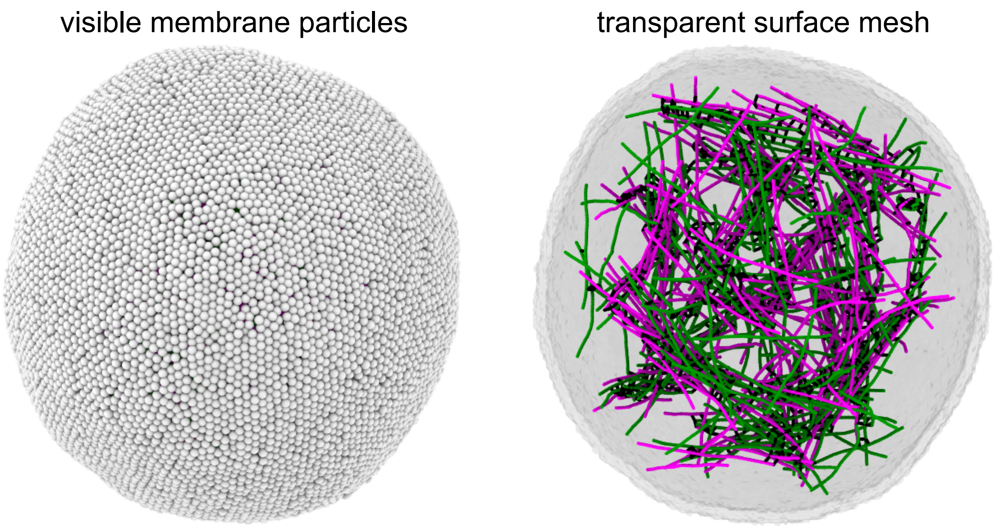


**Figure S12. Visualization of a coarse-grained vesicle of radius $R \approx 24\sigma$ encapsulating a mixed network of crosslinked semiflexible polymers.** (Left) Membrane particles are shown explicitly as spheres, obscuring the vesicle interior. (Right) Membrane rendered as a surface mesh with the half closest to the viewer removed, revealing the internal network architecture.

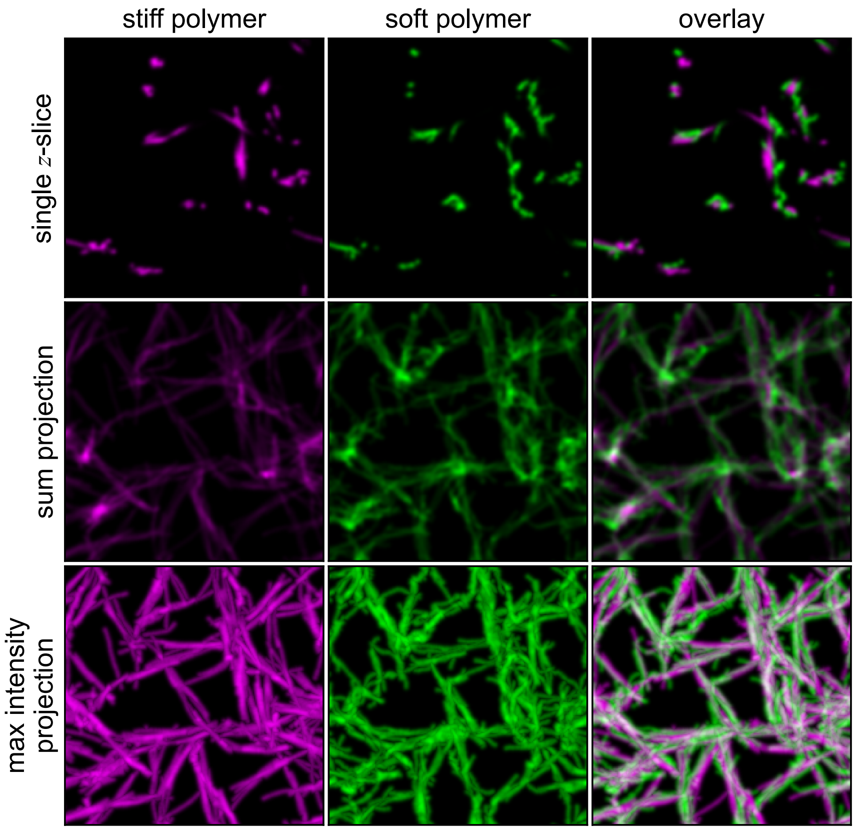


**Figure S13. Visualizing colocalization in bulk network simulations.** Representative artificial fluorescence images for a simulated bulk assembly of crosslinked stiff (magenta) and soft (green) semiflexible polymers are shown alongside an overlay of both channels. The first, second, and third rows shows a single $z$-slice, a sum projection along the $z$-axis, and a maximum intensity projection along the $z$-axis, respectively. Here the bending stiffness ratio is $\kappa_{\text{soft}}/\kappa_{\text{stiff}} = 0.05$, the crosslinker coverage fraction is $\phi = 0.4$, the concentration of each polymer species is $\rho_{\text{soft}} = \rho_{\text{stiff}} = 0.025\rho^{-3}$, and the side length of the periodic simulation domain is $L_{\text{box}} = 60\sigma$.

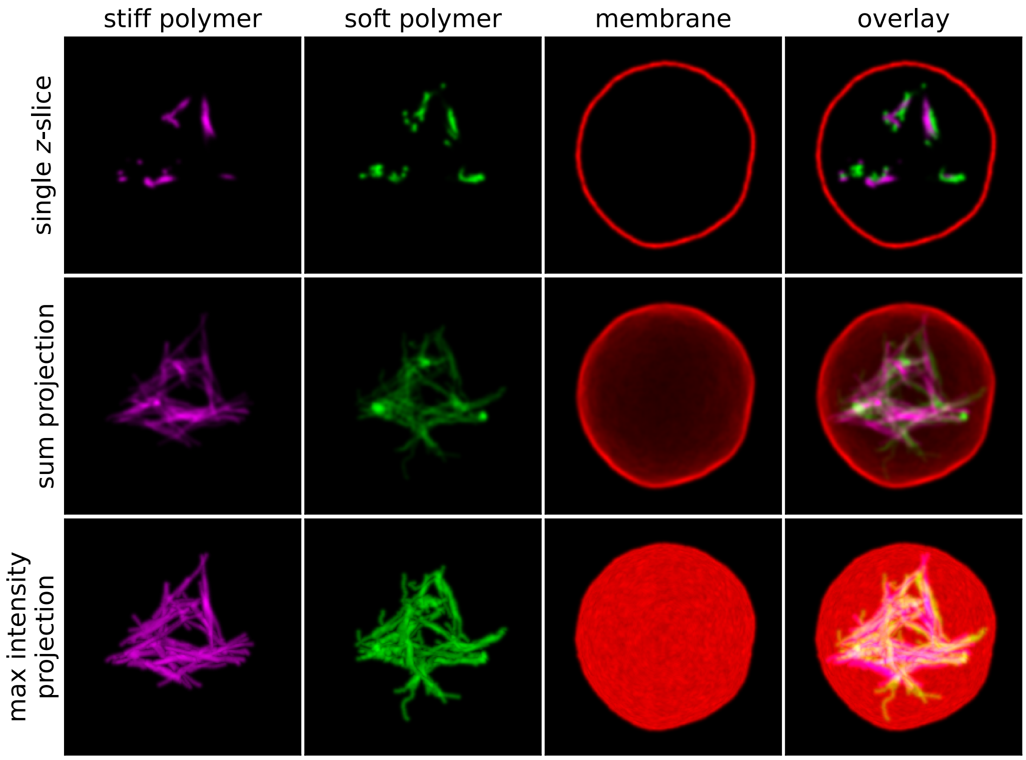


**Figure S14. Visualizing colocalization in vesicle-confined network simulations.** Representative artificial fluorescence images for a simulated crosslinked network of stiff (magenta) and soft (green) semiflexible polymers enclosed in a deformable vesicle (red) are shown alongside an overlay of all three channels. The first, second, and third rows shows a single $z$-slice, a sum projection along the $z$-axis, and a maximum intensity projection along the $z$-axis, respectively. Here the bending stiffness ratio is $\kappa_{\text{soft}}/\kappa_{\text{stiff}} = 0.05$, the crosslinker coverage fraction is $\phi = 0.8$, the concentration of each polymer species is $\rho_{\text{soft}} = \rho_{\text{stiff}} = 0.025\rho^{-3}$, and the average vesicle radius is $R = 27\sigma$. The width of the field of view is $60\sigma$.

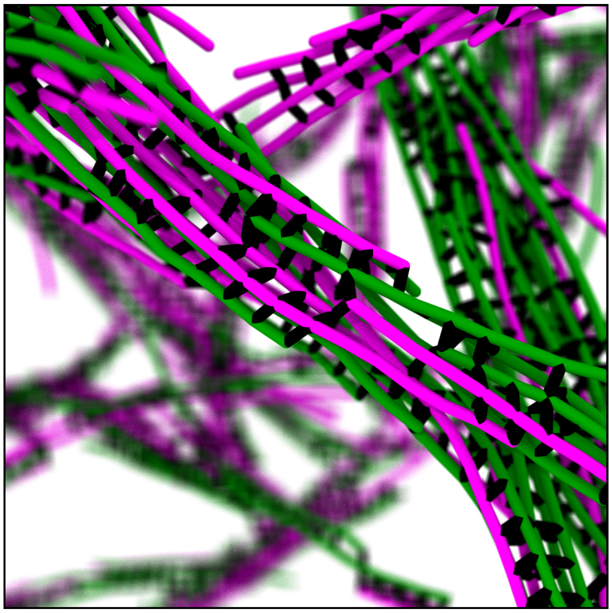

**Figure S15. Cooperative bundling.** When the bending stiffnesses of the two polymer species are similar, we see examples of bundle formation in which entire soft filaments are aligned and crosslinked with stiff filaments. The image focuses on one such bundle in a bulk system. Here, the stiffness ratio of the two polymer species is $\kappa_{\text{soft}}/\kappa_{\text{stiff}} = 0.5$, the crosslinker coverage fraction is $\phi = 0.4$, and the number densities of the two polymer species are $\rho_{\text{soft}} = \rho_{\text{stiff}} = 0.025\sigma^{-3}$.

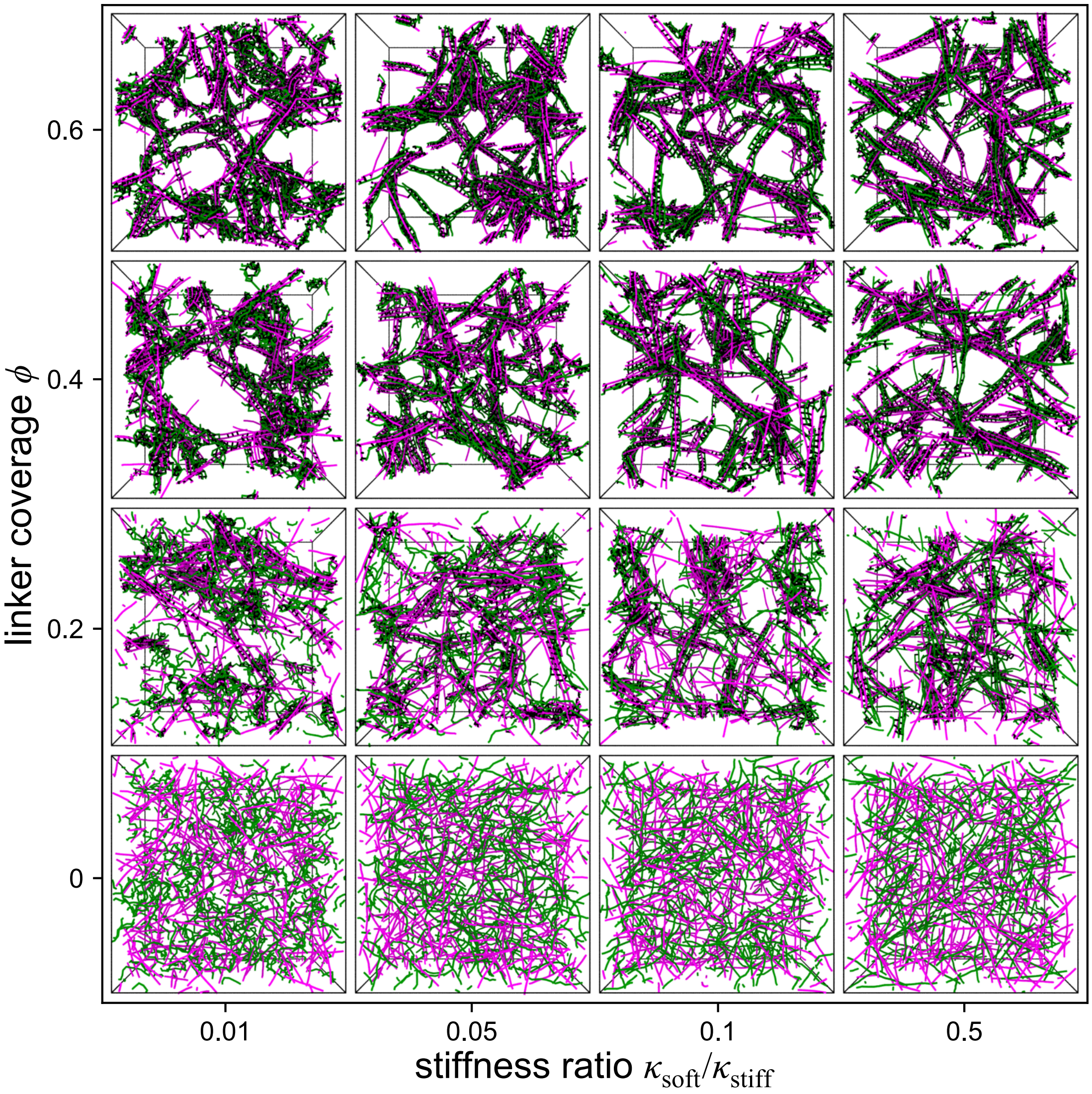


**Figure S16. Bulk assembly of composite networks with varying crosslinker coverage fraction $\phi$ and varying bending stiffness ratio $\kappa_{\text{soft}}/\kappa_{\text{stiff}}$.** Images show stiff (magenta) and soft (green) semiflexible polymers in a periodic simulation domain of side length $L_{\text{box}} = 40\sigma$. Crosslinks are colored black. Each image depicts the final simulation configuration of a representative sample. For these simulations, the particle number density of both species is $\rho_{\text{soft}} = \rho_{\text{stiff}} = 0.05\sigma^{-3}$ and the filament contour length is $L_{\text{soft}} = L_{\text{stiff}} = 14\sigma$. As in the vesicle-confined simulations, increased bending stiffness ratio $\kappa_{\text{soft}}/\kappa_{\text{stiff}}$ and increased crosslinker coverage $\phi$ lead to the formation of aligned bundles consisting of both polymer species.

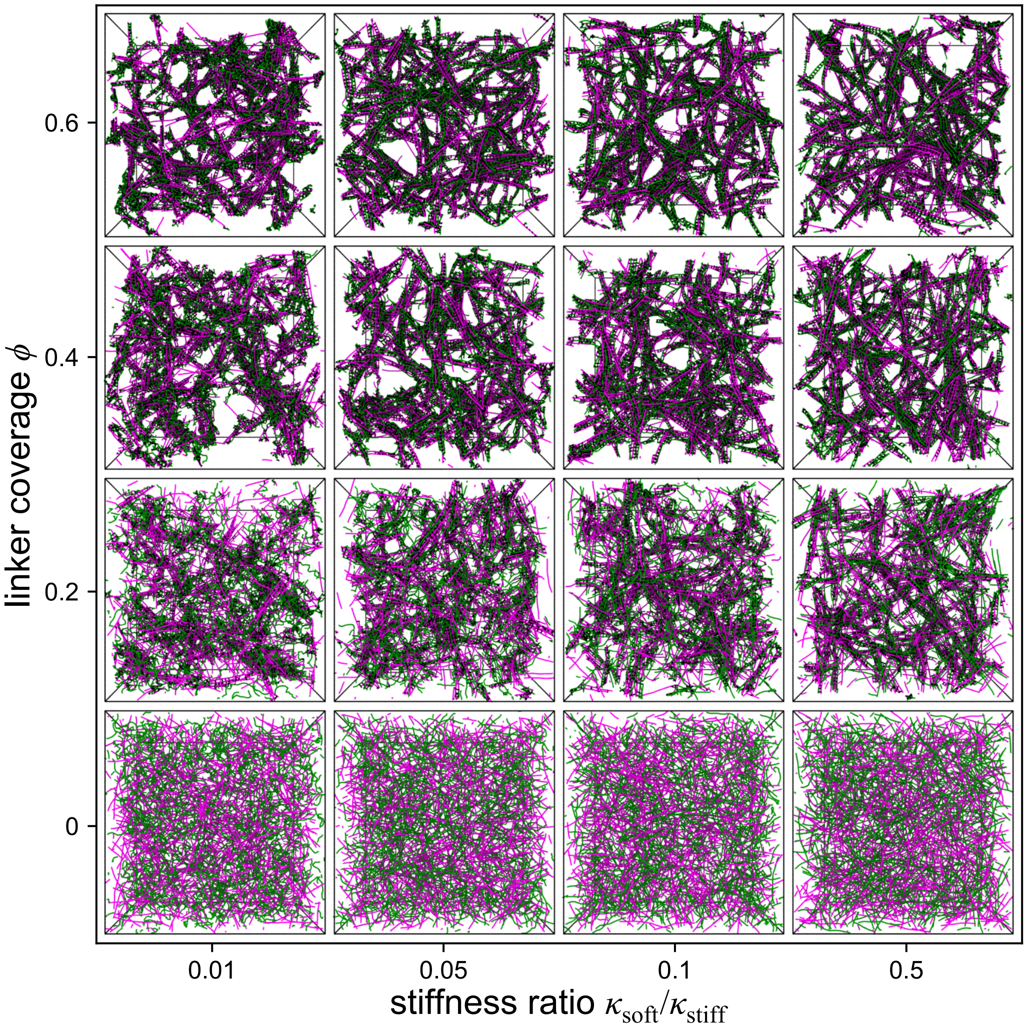


**Figure S17. Bulk assembly of large composite networks with varying crosslinker coverage fraction $\phi$ and varying bending stiffness ratio $\kappa_{\text{soft}}/\kappa_{\text{stiff}}$, for polymer concentration $\rho_{\text{soft}} = \rho_{\text{stiff}} = 0.05\sigma^{-3}$.** Images show stiff (magenta) and soft (green) semiflexible polymers in a periodic simulation domain of side length $L_{\text{box}} = 60\sigma$. Crosslinks are colored black. Each image depicts the final simulation configuration of a representative sample. For these simulations, the filament contour length is $L_{\text{soft}} = L_{\text{stiff}} = 14\sigma$.

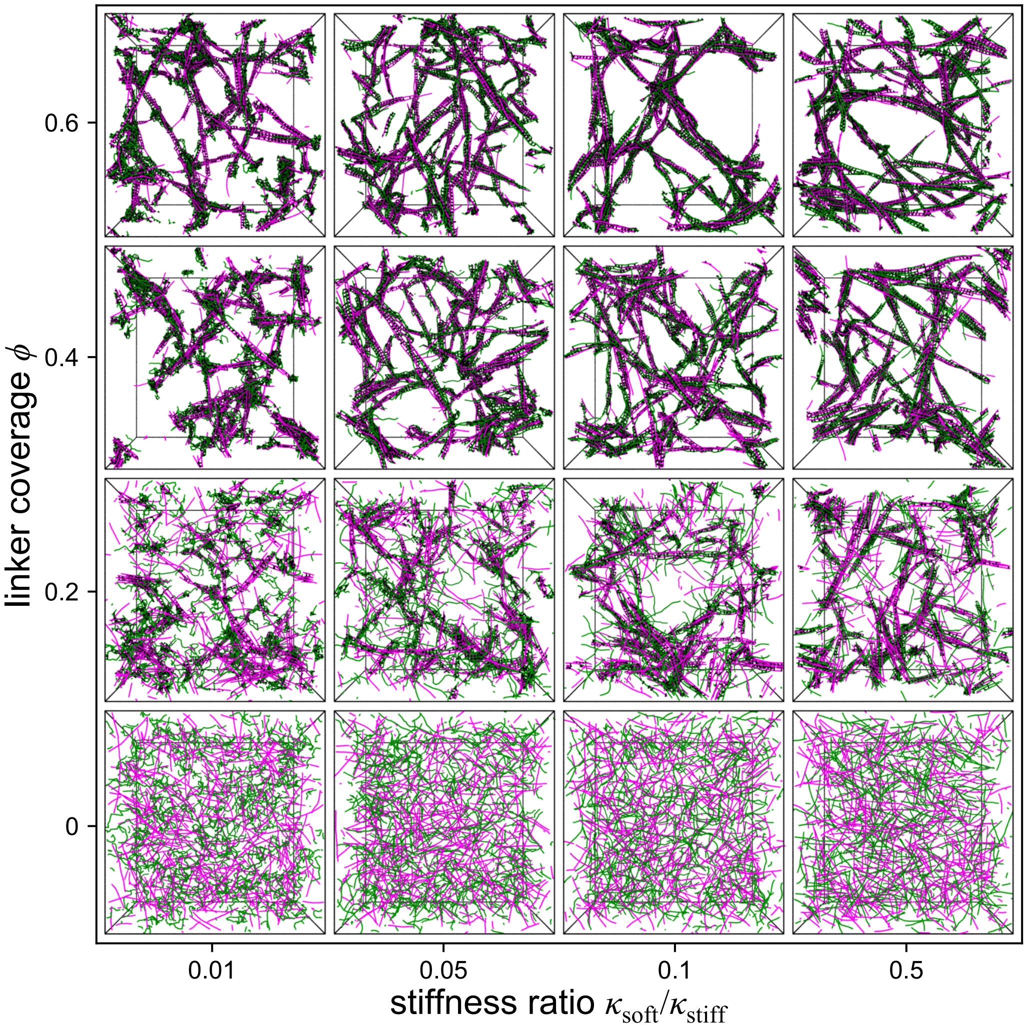


**Figure S18. Bulk assembly of large composite networks with varying crosslinker coverage fraction $\phi$ and varying bending stiffness ratio $\kappa_{\text{soft}}/\kappa_{\text{stiff}}$, for polymer concentration $\boldsymbol{\rho_{\text{soft}} = \rho_{\text{stiff}} = 0.025\sigma^{-3}}$.** Images show stiff (magenta) and soft (green) semiflexible polymers in a periodic simulation domain of side length $L_{\text{box}} = 60\sigma$. Crosslinks are colored black. Each image depicts the final simulation configuration of a representative sample. For these simulations, the filament contour length is $L_{\text{soft}} = L_{\text{stiff}} = 14\sigma$.

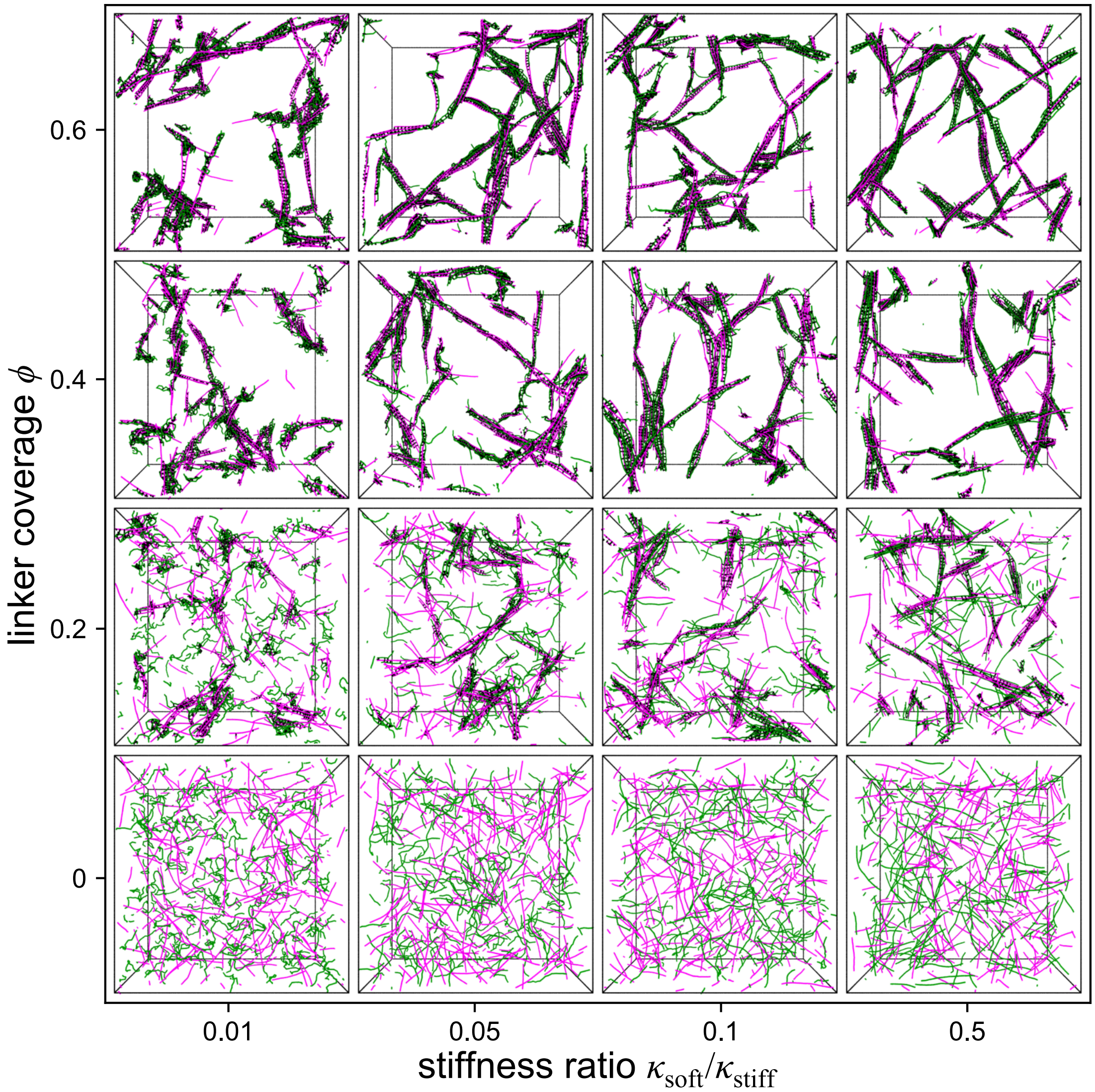


**Figure S19. Bulk assembly of large composite networks with varying crosslinker coverage fraction $\phi$ and varying bending stiffness ratio $\kappa_{\mathrm{soft}}/\kappa_{\mathrm{stiff}}$, for polymer concentration $\boldsymbol{\rho}_{\mathbf{soft}} = \boldsymbol{\rho}_{\mathbf{stiff}} = \mathbf{0.0125}\boldsymbol{\sigma}^{-\mathbf{3}}$.** Images show stiff (magenta) and soft (green) semiflexible polymers in a periodic simulation domain of side length $L_{\mathrm{box}} = 80\sigma$. Crosslinks are colored black. Each image depicts the final simulation configuration of a representative sample. For these simulations, the filament contour length is $L_{\mathrm{soft}} = L_{\mathrm{stiff}} = 14\sigma$.

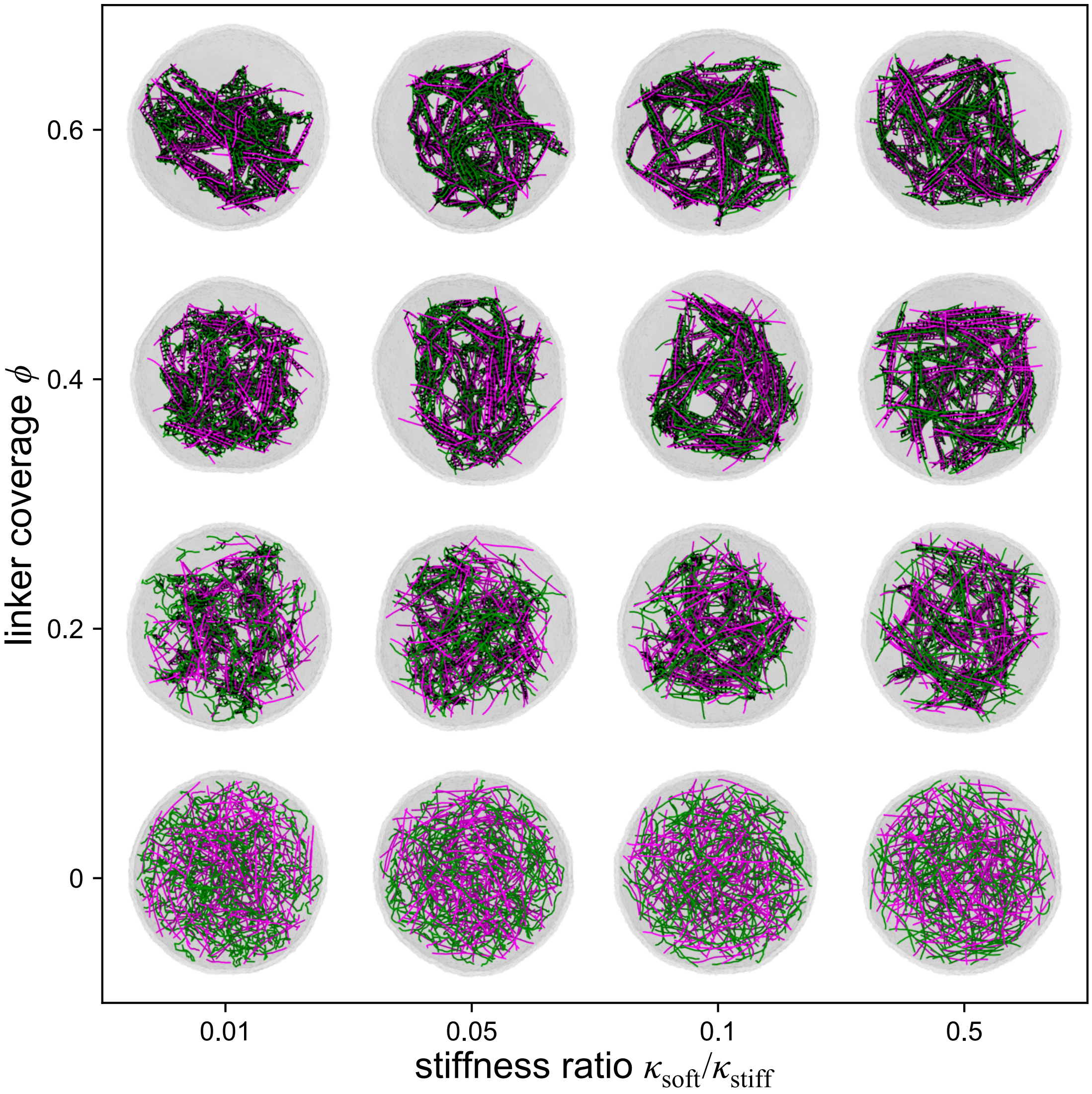


**Figure S20. Confined assembly of composite networks with varying crosslinker coverage fraction $\phi$ and varying bending stiffness ratio $\kappa_{\text{soft}}/\kappa_{\text{stiff}}$.** Images show stiff (magenta) and soft (green) semiflexible polymers encapsulated in a deformable vesicle, rendered as a semi-transparent surface. Crosslinks are colored black. Each image depicts the final simulation configuration of a representative sample. For these simulations, the particle number density of both species is $\rho_{\text{soft}} = \rho_{\text{stiff}} = 0.05\sigma^{-3}$, the filament contour length is $L_{\text{soft}} = L_{\text{stiff}} = 14\sigma$, and the vesicle radius is $R = 27\sigma$. Increased bending stiffness ratio $\kappa_{\text{soft}}/\kappa_{\text{stiff}}$ and increased crosslinker coverage fraction $\phi$ lead to the formation of aligned bundles consisting of both polymer species.

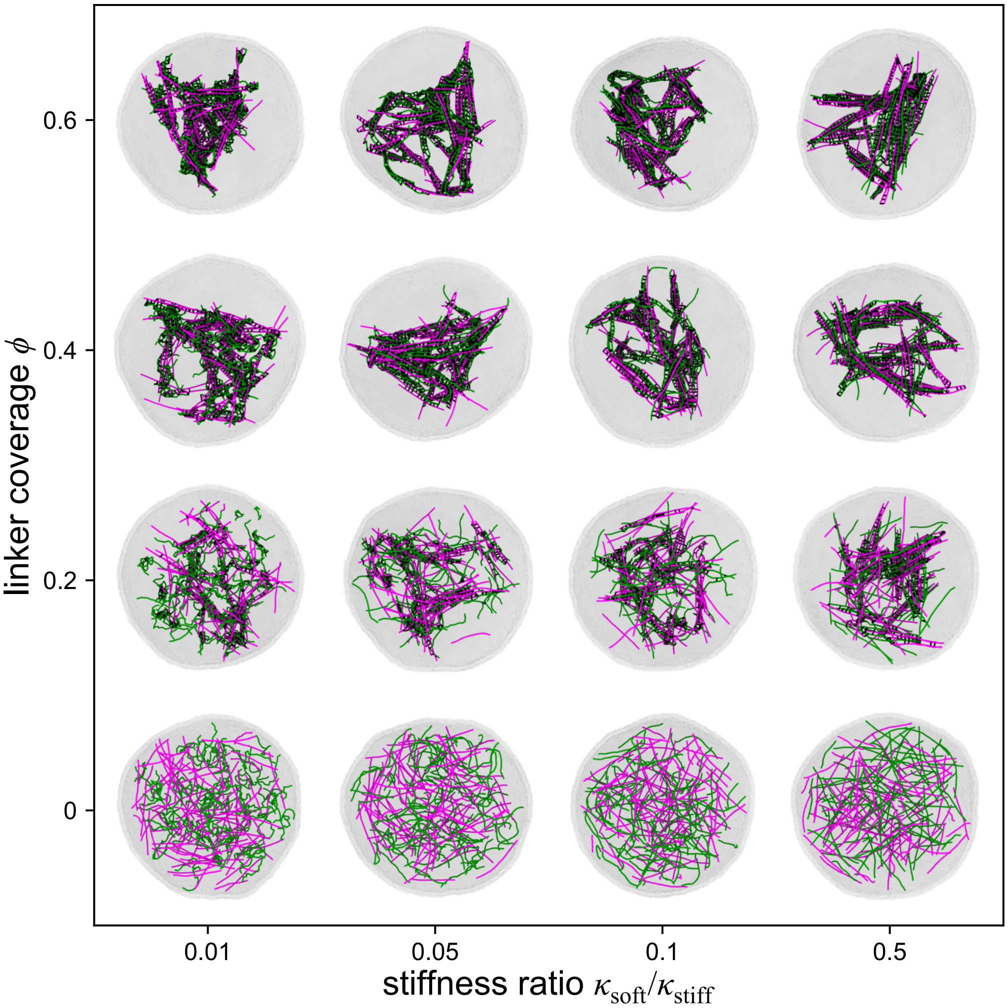


**Figure S21. Confined assembly of composite networks with varying crosslinker coverage fraction $\phi$ and varying bending stiffness ratio $\kappa_{\text{soft}}/\kappa_{\text{stiff}}$ for polymer concentration $\rho_{\text{soft}} = \rho_{\text{stiff}} = 0.025\sigma^{-3}$ .** Images show stiff (magenta) and soft (green) semiflexible polymers encapsulated in a deformable vesicle, rendered as a semi-transparent surface. Crosslinks are colored black. Each image depicts the final simulation configuration of a representative sample. For these simulations, the filament contour length is $L_{\text{soft}} = L_{\text{stiff}} = 14\sigma$ and the vesicle radius is $R = 27\sigma$. As is observed in the higher concentration case, increased bending stiffness ratio $\kappa_{\text{soft}}/\kappa_{\text{stiff}}$ and increased crosslinker coverage fraction $\phi$ lead to the formation of aligned bundles consisting of both polymer species.

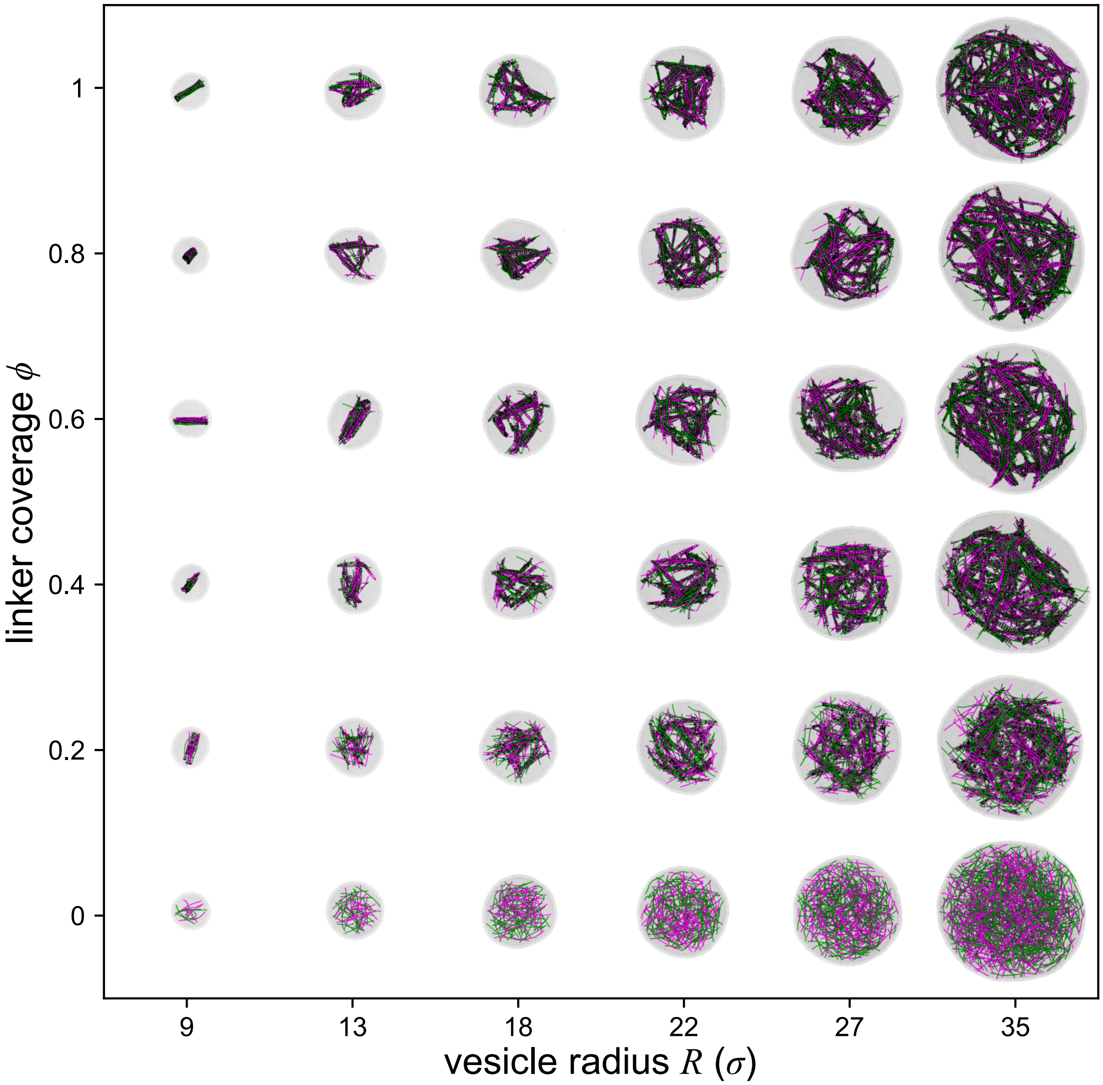


**Figure S22. Confined assembly of composite networks with varying crosslinker coverage fraction $\phi$ within vesicles of varying radius $R$ for polymer concentration $\rho_{\text{soft}} = \rho_{\text{stiff}} = 0.05\sigma^{-3}$.** Images show stiff (magenta) and soft (green) semiflexible polymers encapsulated in a deformable vesicle, rendered as a semi-transparent surface. Crosslinks are colored black. Each image depicts the final simulation configuration of a representative sample. For these simulations, the filament contour length is $L_{\text{soft}} = L_{\text{stiff}} = 14\sigma$ and the bending stiffness ratio is $\kappa_{\text{soft}}/\kappa_{\text{stiff}} = 0.5$. Increased confinement (smaller $R$) and increased crosslinker coverage fraction $\phi$ lead to a shift from extended internal network architectures to more compact, bundled architectures.

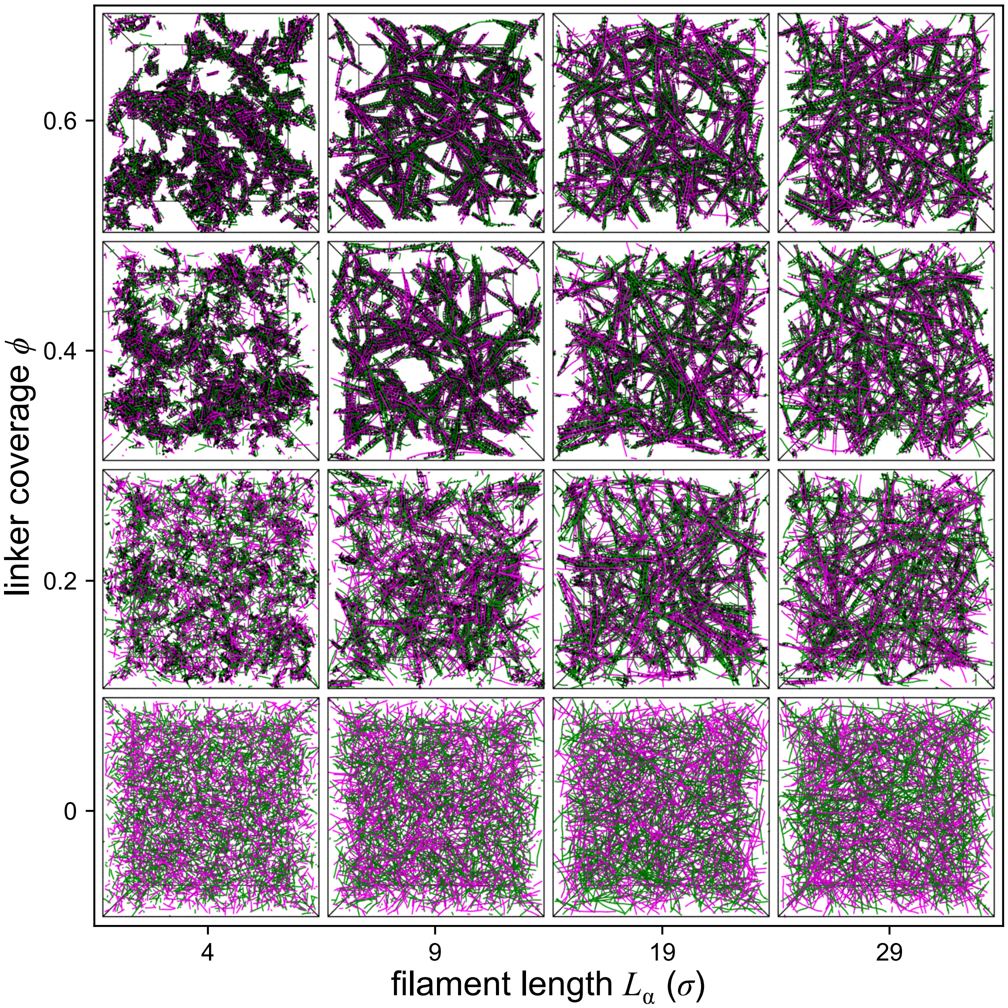


**Figure S23. Bulk assembly of large composite networks with varying crosslinker coverage fraction $\phi$ and varying filament length $L_\alpha$.** Images show stiff (magenta) and soft (green) semiflexible polymers in a periodic simulation domain of side length $L_{\text{box}} = 60\sigma$. Crosslinks are colored black. Each image depicts the final simulation configuration of a representative sample. For these simulations, the bending stiffness ratio is $\kappa_{\text{soft}}/\kappa_{\text{stiff}} = 0.5$, and the polymer concentration is $\rho_{\text{soft}} = \rho_{\text{stiff}} = 0.05\sigma^{-3}$.

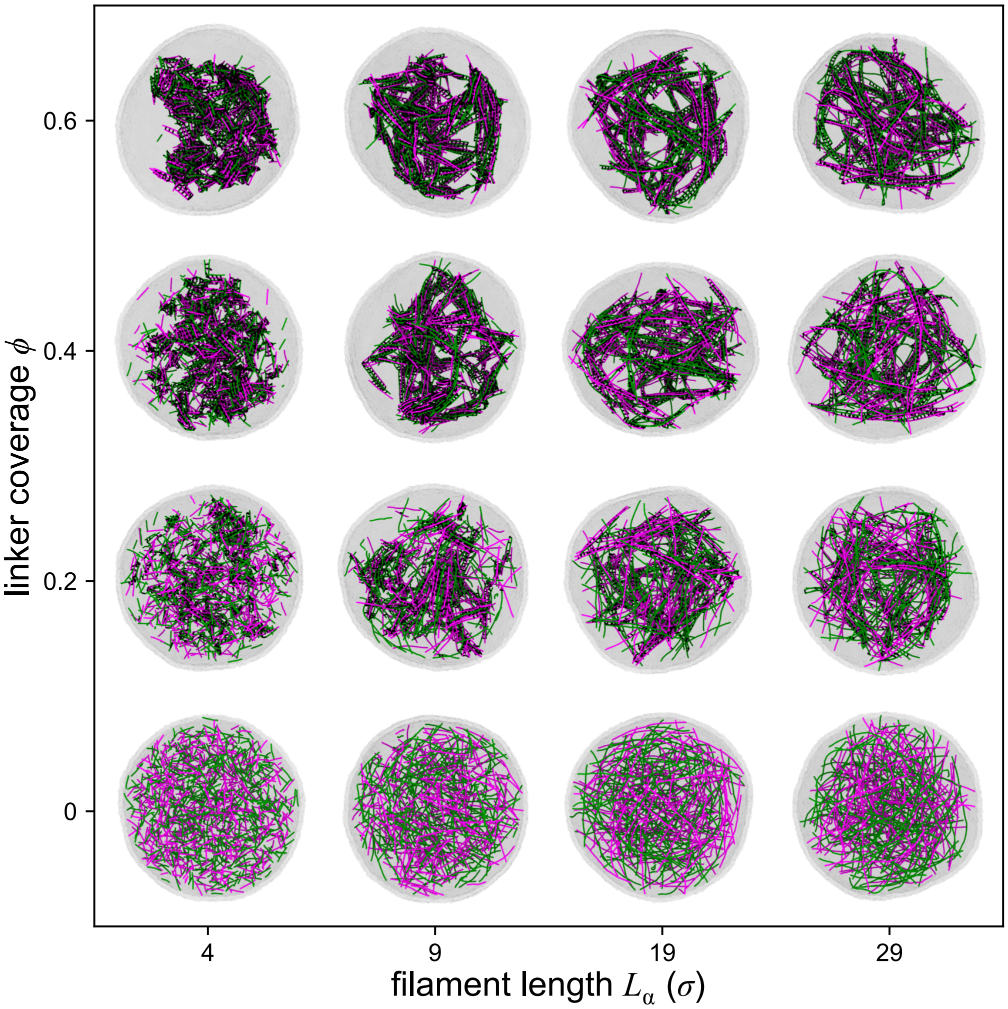


**Figure S24. Confined assembly of composite networks with varying crosslinker coverage fraction $\phi$ and varying filament length $L_\alpha$.** Images show stiff (magenta) and soft (green) semiflexible polymers encapsulated in a deformable vesicle, rendered as a semi-transparent surface. Crosslinks are colored black. Each image depicts the final simulation configuration of a representative sample. For these simulations, the vesicle radius is $R = 27\sigma$, the bending stiffness ratio is $\kappa_{\text{soft}}/\kappa_{\text{stiff}} = 0.5$, and the polymer concentration is $\rho_{\text{soft}} = \rho_{\text{stiff}} = 0.05\sigma^{-3}$.

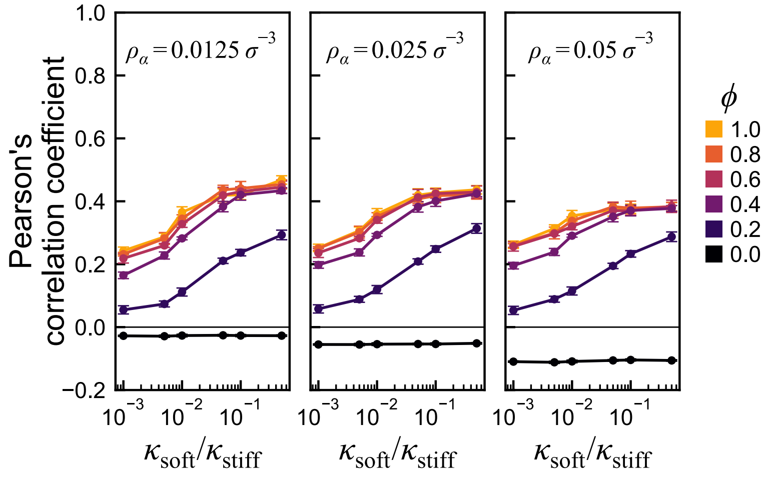


**Figure S25. Comparison of stiff-soft polymer colocalization measures in bulk systems with varied polymer concentrations**. Pearson's correlation coefficient is plotted as a function of the stiffness ratio $\kappa_{\text{soft}}/\kappa_{\text{stiff}}$ with varied crosslinker coverage fraction $\phi$ in periodic bulk systems with $L_{\text{box}} = 60\sigma$ with varying values of the polymer concentration $\rho_\alpha$ , in which $\alpha \in \{\text{stiff}, \text{soft}\}$ corresponds to the polymer species. Note that reducing the polymer concentration $\rho_\alpha$ weakly increases colocalization overall and mitigates the decrease in colocalization with increasing $\kappa_{\text{soft}}/\kappa_{\text{stiff}}$ seen for large values of $\phi$.

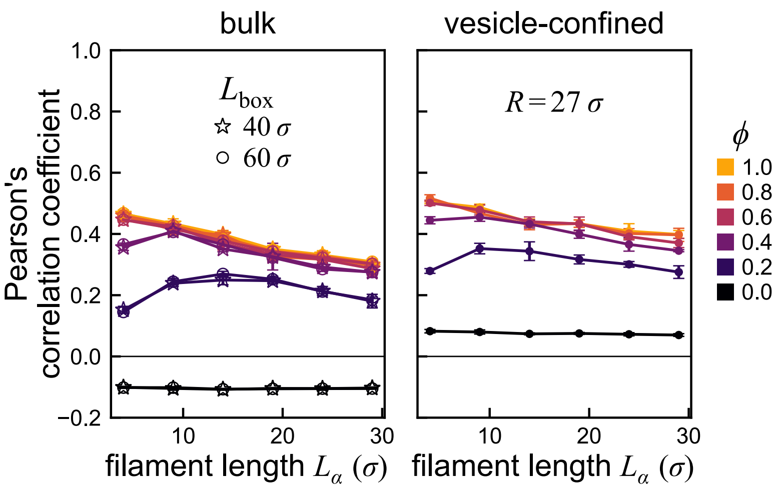


**Figure S26. Comparison of stiff-soft polymer colocalization measures in bulk and vesicle-confined systems with varied filament length $L_\alpha$.** Pearson's correlation coefficient is plotted as a function of the filament length $L_\alpha$, in which $\alpha \in$ {stiff, soft}, for filaments with varied crosslinker coverage fraction $\phi$ in (left panel) periodic bulk networks of two system sizes and (right panel) networks confined in vesicles of radius $R = 27\sigma$. For these simulations, $\kappa_{\text{soft}}/\kappa_{\text{stiff}} = 0.5$.

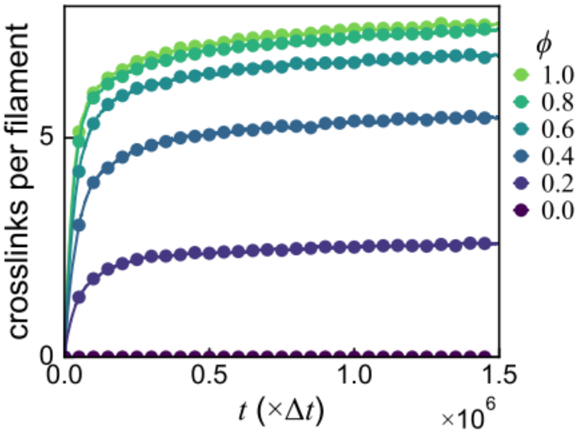


**Figure S27. Time evolution of the average number of crosslinks per filament at different coverage fractions ($\phi$)**, shown for a representative set of simulation parameters.

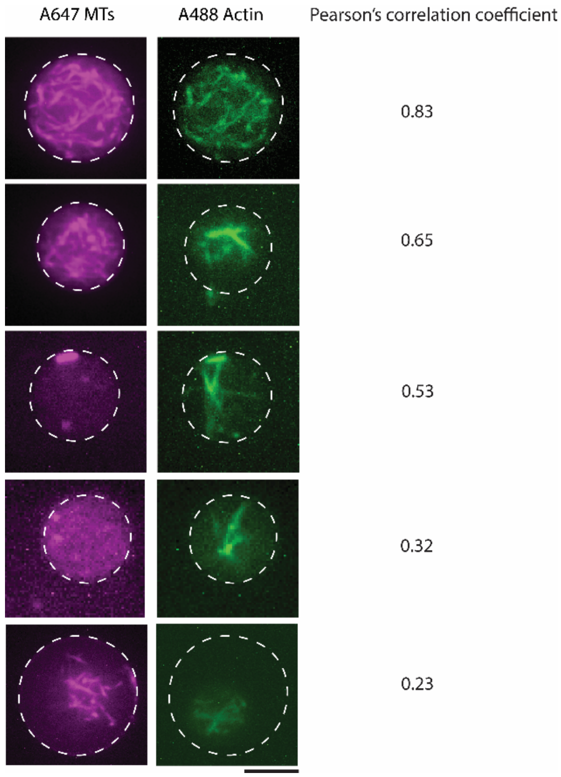


**Figure S28. Representative confocal fluorescence microscopy images** illustrating MT-actin colocalization corresponding to varying Pearson correlation coefficients. Scale bar: 10 μm.

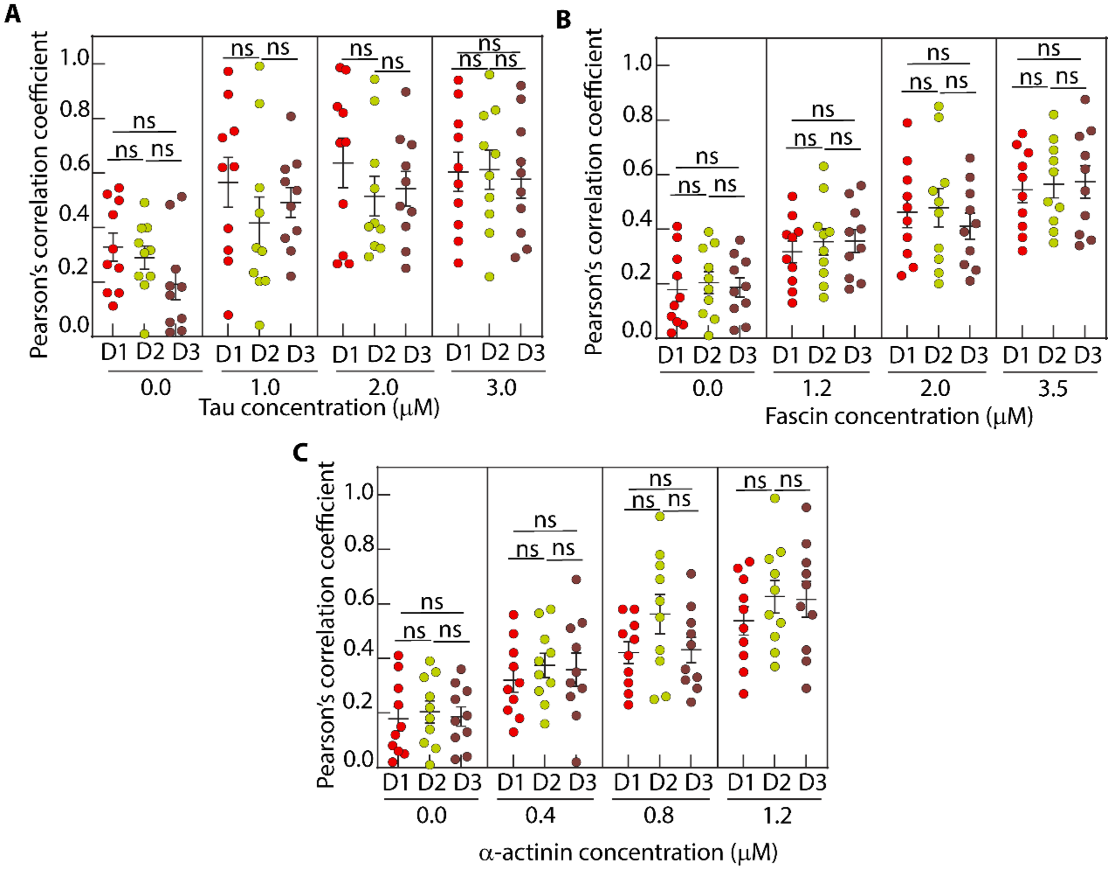


**Figure S29. Distribution of Pearson's correlation coefficients obtained from three independent experimental days (D1, D2, and D3) at varying concentrations** of (A) Tau (with fascin fixed at 1.2 μM), (B) Fascin (with tau fixed at 2.0 μM), and (C) α-actinin (with tau fixed at 2.0 μM). ns indicates non-significant differences.

### Supplementary Table

#### Table S1. List of p-values

| Figure | Condition | Comparison values | p-values |
|---|---|---|---|
| **2-E** | Tau concentration | 0.0 μM (fascin -) vs 0.0 μM (fascin+) | 0.0857 |
| | | 0.0 μM (Fascin +) vs 1.0 μM (Fascin -) | $2.73\times10^{-4}$ |
| | | 1.0 μM (Fascin -) vs 1.0 μM (Fascin +) | 0.0110 |
| | | 1.0 μM (Fascin +) vs 2.0 μM (Fascin +) | $3.30\times10^{-11}$ |
| | | 0.0 μM (Fascin +) vs 1.0 μM (Fascin +) | $7.20\times10^{-11}$ |
| | | 1.0 μM (Fascin -) vs 2.0 μM (Fascin +) | $<10^{-12}$ |
| | | 0.0 μM (Fascin +) vs 2.0 μM (Fascin +) | $<10^{-12}$ |
| **2-G** | Tau concentration | 0.0 μM (-α-actinin) vs 0.0 μM (+α-actinin) | 0.9837 |
| | | 0.0 μM Tau (+α-actinin) vs 1.0 μM Tau (+α-actinin) | $<10^{-12}$ |
| | | 1.0 μM Tau (+α-actinin) vs 2.0 μM Tau (+α-actinin) | $<10^{-12}$ |
| | | 0.0 μM (-α-actinin) vs 1.0 μM (+α-actinin) | 0.0539 |
| | | 0.0 μM Tau (+α-actinin) vs 2.0 μM Tau (+α-actinin) | 0.0043 |
| **3-D** | Tau concentration | 0.0 μM vs 1.0 μM | $1.21\times10^{-3}$ |
| | | 1.0 μM vs 2.0 μM | 0.7506 |
| | | 2.0 μM vs 3.0 μM | 0.9934 |
| | | 0.0 μM vs 2.0 μM | $7.76\times10^{-7}$ |
| | | 0.0 μM vs 3.0 μM | $6.03\times10^{-8}$ |
| | | 1.0 μM vs 3.0 μM | 0.3435 |
| **3E** | Tau concentration | 0.0 μM (7-14 μm vs. 14-21 μm) | 0.3362 |
| | | 0.0 μM (14-21 μm vs. >21μm) | 0.0343 |
| | | 0.0 μM (7-14 μm vs. >21μm) | 0.0186 |
| | | 1.0 μM (7-14 μm vs. 14-21 μm) | 0.3209 |
| | | 1.0 μM (14-21 μm vs. >21μm) | 0.0490 |
| | | 1.0 μM (7-14 μm vs. >21μm) | 0.8391 |

| | | | |
|---|---|---|---|
| | | 2.0 μM (7-14 μm vs. 14-21 μm) | 0.0362 |
| | | 2.0 μM (14-21 μm vs. >21μm) | 0.0271 |
| | | 2.0 μM (7-14 μm vs. >21μm) | 0.8046 |
| | | 3.0 μM (7-14 μm vs. 14-21 μm) | 0.6249 |
| | | 3.0 μM (14-21 μm vs. >21μm) | 0.9257 |
| | | 3.0 μM (7-14 μm vs. >21μm) | 0.5492 |
| **4B** | Fascin concentration | 0.0 μM vs 1.2 μM | $9.99\times10^{-4}$ |
| | | 1.2 μM vs 2.0 μM | 0.0410 |
| | | 2.0 μM vs 3.5 μM | 0.0351 |
| | | 0.0 μM vs 2.0 μM | $6.34\times10^{-9}$ |
| | | 0.0 μM vs 3.5 μM | $<10^{-12}$ |
| **4C** | Fascin concentration | 0.0 μM (7-14 μm vs. 14-21 μm) | 0.5978 |
| | | 0.0 μM (14-21 μm vs. >21μm) | 0.4383 |
| | | 0.0 μM (7-14 μm vs. >21μm) | 0.1721 |
| | | 1.2 μM (7-14 μm vs. 14-21 μm) | 0.8209 |
| | | 1.2 μM (14-21 μm vs. >21μm) | 0.5067 |
| | | 1.2 μM (7-14 μm vs. >21μm) | 0.7724 |
| | | 2.0 μM (7-14 μm vs. 14-21 μm) | 0.0447 |
| | | 2.0 μM (14-21 μm vs. >21μm) | 0.3596 |
| | | 2.0 μM (7-14 μm vs. >21μm) | 0.8011 |
| | | 3.5 μM (7-14 μm vs. 14-21 μm) | 0.8219 |
| | | 3.5 μM (14-21 μm vs. >21μm) | 0.1570 |
| | | 3.5 μM (7-14 μm vs. >21μm) | 0.2889 |
| **4E** | α-actinin concentration | 0.0 μM vs 0.4 μM | $1.12\times10^{-3}$ |
| | | 0.4 μM vs 0.8 μM | 0.0296 |
| | | 0.8 μM vs 1.2 μM | 0.2585 |
| | | 0.0 μM vs 0.8 μM | $4.32\times10^{-9}$ |
| | | 0.0 μM vs 1.2 μM | $<10^{-12}$ |
| **4F** | α-actinin concentration | 0.0 μM (7-14 μm vs. 14-21 μm) | 0.5978 |
| | | 0.0 μM (14-21 μm vs. >21μm) | 0.4383 |
| | | 0.0 μM (7-14 μm vs. >21μm) | 0.1721 |
| | | 0.4 μM (7-14 μm vs. 14-21 μm) | 0.2169 |

| | | | |
|---|---|---|---|
| | | 0.4 μM (14-21 μm vs. >21μm) | 0.0024 |
| | | 0.4 μM (7-14 μm vs. >21μm) | 0.0726 |
| | | 0.8 μM (7-14 μm vs. 14-21 μm) | 0.0451 |
| | | 0.8 μM (14-21 μm vs. >21μm) | 0.0423 |
| | | 0.8 μM (7-14 μm vs. >21μm) | 0.7234 |
| | | 1.2 μM (7-14 μm vs. 14-21 μm) | 0.0711 |
| | | 1.2 μM (14-21 μm vs. >21μm) | 0.0712 |
| | | 1.2 μM (7-14 μm vs. >21μm) | $3.41\times10^{-3}$ |
| **S7** | Pearson's coefficient | Control/-MC vs. Control/+MC | 0.9729 |
| | | Fascin/-MC vs Fascin/+MC | 0.1794 |
| | | α-actinin /-MC vs α-actinin /+MC | 0.7709 |
| **S8** | Pearson's coefficient | TIRF vs confocal | 0.0726 |
| **S29** | Pearson's coefficient | Tau 0 μM (D1 vs. D2) | >0.9999 |
| | | Tau 0 μM (D2 vs. D3) | 0.9999 |
| | | Tau 0 μM (D1 vs. D3) | 0.9999 |
| | | Tau 1 μM (D1 vs. D2) | 0.9999 |
| | | Tau 1 μM (D2 vs. D3) | >0.9999 |
| | | Tau 1 μM (D1 vs. D3) | >0.9999 |
| | | Tau 2 μM (D1 vs. D2) | 0.9999 |
| | | Tau 2 μM (D2 vs. D3) | >0.9999 |
| | | Tau 2 μM (D1 vs. D3) | >0.9999 |
| | | Tau 3 μM (D1 vs. D2) | >0.9999 |
| | | Tau 3 μM (D2 vs. D3) | >0.9999 |
| | | Tau 3 μM (D1 vs. D3) | >0.9999 |
| | | Fascin 0 μM (D1 vs. D2) | >0.9999 |
| | | Fascin 0 μM (D2 vs. D3) | >0.9999 |
| | | Fascin 0 μM (D1 vs. D3) | >0.9999 |
| | | Fascin 1.2 μM (D1 vs. D2) | >0.9999 |
| | | Fascin 1.2 μM (D2 vs. D3) | >0.9999 |
| | | Fascin 1.2 μM (D1 vs. D3) | >0.9999 |
| | | Fascin 2.0 μM (D1 vs. D2) | >0.9999 |
| | | Fascin 2.0 μM (D2 vs. D3) | 0.9999 |
| | | Fascin 2.0 μM (D1 vs. D3) | >0.9999 |

| | | | |
|---|---|---|---|
| | | Fascin 3.5 μM (D1 vs. D2) | >0.9999 |
| | | Fascin 3.5μM (D2 vs. D3) | >0.9999 |
| | | Fascin 3.5 μM (D1 vs. D3) | >0.9999 |
| | | α-actinin 0 μM (D1 vs. D2) | >0.9999 |
| | | α-actinin 0 μM (D2 vs. D3) | >0.9999 |
| | | α-actinin 0 μM (D1 vs. D3) | >0.9999 |
| | | α-actinin 0.4 μM (D1 vs. D2) | >0.9999 |
| | | α-actinin 0.4 μM (D2 vs. D3) | >0.9999 |
| | | α-actinin 0.4 μM (D1 vs. D3) | >0.9999 |
| | | α-actinin 0.8 μM (D1 vs. D2) | 0.9772 |
| | | α-actinin 0.8 μM (D2 vs. D3) | 0.9942 |
| | | α-actinin 0.8 μM (D1 vs. D3) | >0.9999 |
| | | α-actinin 1.2 μM (D1 vs. D2) | 0.9999 |
| | | α-actinin 1.2 μM (D2 vs. D3) | 0.9999 |
| | | α-actinin 1.2 μM (D1 vs. D3) | >0.9999 |

**Table S2. Simulation parameters.** Physical units are specified in terms of characteristic energy $\varepsilon$, mass $m$, and distance $\sigma$. Quantities for which the value entry is "varied" are specified in the text.

| symbol | value | description |
|---|---|---|
| ε | 1 | unit energy |
| $m$ | 1 | unit mass |
| σ | 1 | unit length |
| $\tau_0$ | $\sigma\sqrt{m/\varepsilon}$ | characteristic timescale |
| $k_\mathrm{B}T$ | 1.0 ε | thermal energy scale |
| $\gamma^{-1}$ | $1.0\ \tau_0$ | damping coefficient |
| $\Delta t$ | $0.0025\ \tau_0$ | simulation timestep |
| $L_\mathrm{box}$ | Varied | side length of periodic simulation box (cubic) |
| $\sigma_\mathrm{m}$ | $1.0\ \sigma$ | membrane particle diameter |
| $\varepsilon_\mathrm{m}$ | $4.34\ \varepsilon$ | energy scale for membrane-membrane particle interaction [2] |
| $r_\mathrm{min}$ | $2^{1/6}\ \sigma_\mathrm{m}$ | distance corresponding to minimum in $U_\mathrm{mem}$ [2] |
| $r_c^m$ | $2.6\ \sigma_\mathrm{m}$ | membrane-membrane particle interaction cutoff distance [2] |
| $\zeta$ | 2.5 | exponent controlling in-plane diffusivity of membrane particles [2] |
| $\mu$ | 3 | "aligning stiffness" in $U_\mathrm{mem}$; controls bending rigidity [2] |
| $\theta_0$ | 0 | membrane spontaneous curvature |
| $\sigma_\mathrm{p}$ | $0.8\ \sigma$ | polymer particle diameter |
| $\varepsilon_\mathrm{p}$ | $1.0\ \varepsilon$ | characteristic energy scale for polymer-polymer repulsion |
| $l_0$ | $1.0\ \sigma$ | polymer spring rest length |
| $\theta_0^\mathrm{p}$ | 0 | polymer spontaneous curvature |
| $\varepsilon_\mathrm{mp}$ | $4.34\ \varepsilon$ | energy scale for membrane-polymer interaction [2] |
| $\tau_\mathrm{equil}$ | $2 \times 10^5 \Delta t$ | duration of equilibration stage |
| $\tau_\mathrm{main}$ | $1.5 \times 10^6 \Delta t$ | duration of main (crosslinking) stage |
| $k_\mathrm{stretch}^\mathrm{stiff}$ | $300\ \varepsilon/\sigma^2$ | stretching constant for stiff polymers |
| $k_\mathrm{stretch}^\mathrm{soft}$ | $300\ \varepsilon/\sigma^2$ | stretching constant for soft polymers |
| $k_\mathrm{stretch}^\mathrm{cl}$ | $50\ \varepsilon/\sigma^2$ | stretching constant for crosslinks |
| $k_\mathrm{bend}^\mathrm{stiff}$ | $100\ \varepsilon$ | bending constant for stiff polymers |
| $k_\mathrm{bend}^\mathrm{soft}$ | varied | bending constant for soft polymers |

| | | |
|---|---|---|
| $n_{\text{stiff}}$ | 15 | number of particles per stiff polymer |
| $n_{\text{soft}}$ | 15 | number of particles per soft polymer |
| $\rho_{\text{stiff}}$ | $0.05\ \sigma^{-3}$ | number density of stiff polymer particles |
| $\rho_{\text{soft}}$ | $0.05\ \sigma^{-3}$ | number density of soft polymer particles |
| $\phi_{\text{stiff}}$ | varied | crosslinker coverage fraction for stiff polymer sites |
| $\phi_{\text{soft}}$ | varied | crosslinker coverage fraction for soft polymer sites |
| $p_{\text{cl}}^{\text{soft,stiff}}$ | $0.05$ | probability of soft-stiff crosslink formation during a single timestep |
| $p_{\text{cl}}^{\text{stiff,stiff}}$ | $0.05$ | probability of stiff-stiff crosslink formation during a single timestep |
| $p_{\text{cl}}^{\text{soft,soft}}$ | $0.05$ | probability of soft-soft crosslink formation during a single timestep |
| $p_{\text{cl}}^{\text{break}}$ | $10^{-4}$ | probability of crosslink breakage during a single timestep |
| $l_0^{\text{cl}}$ | $0.8\ \sigma$ | rest length for crosslinks |
